# A minimal physical model for curvotaxis driven by curved protein complexes at the cell’s leading edge

**DOI:** 10.1101/2023.04.19.537490

**Authors:** Raj Kumar Sadhu, Marine Luciano, Wang Xi, Cristina Martinez-Torres, Marcel Schröder, Christoph Blum, Marco Tarantola, Samo Penič, Aleš Iglič, Carsten Beta, Oliver Steinbock, Eberhard Bodenschatz, Benoît Ladoux, Sylvain Gabriele, Nir S. Gov

## Abstract

Cells often migrate on curved surfaces inside the body, such as curved tissues, blood vessels or highly curved protrusions of other cells. Recent *in-vitro* experiments provide clear evidence that motile cells are affected by the curvature of the substrate on which they migrate, preferring certain curvatures to others, termed “curvotaxis”. The origin and underlying mechanism that gives rise to this curvature sensitivity are not well understood. Here, we employ a “minimal cell” model which is composed of a vesicle that contains curved membrane protein complexes, that exert protrusive forces on the membrane (representing the pressure due to actin polymerization). This minimal-cell model gives rise to spontaneous emergence of a motile phenotype, driven by a lamellipodia-like leading edge. By systematically screening the behaviour of this model on different types of curved substrates (sinusoidal, cylinder and tube), we show that minimal ingredients and energy terms capture the experimental data. The model recovers the observed migration on the sinusoidal substrate, where cells move along the grooves (minima), while avoiding motion along the ridges. In addition, the model predicts the tendency of cells to migrate circumferentially on convex substrates and axially on concave ones. Both of these predictions are verified experimentally, on several cell types. Altogether, our results identify the minimization of membrane-substrate adhesion energy and binding energy between the membrane protein complexes as key players of curvotaxis in cell migration.

## I. INTRODUCTION

Cell migration is an important biological process that plays a central role in immune response, wound healing, tissue homeostasis etc [1, 2]. While the environment of a cell *in vivo* is geometrically complex, most of the studies focus on cell spreading and migration on flat substrates [3–5]. Previous studies on 2D patterned flat surfaces have shown that cells adapt their shape and their internal cytoskeleton to these 2D geometries [6–8]. However, eukaryote cells, adhering and migrating on a solid substrate, are observed to also interact with the topography of the substrate and modify their motility [9, 10]. The alignment and the direction of migration of isolated cells, in response to the topography, crucially depends upon the cell type. For example, fibroblasts are found to align axially on the surface of a cylinder, while epithelial cell align circumferentially [11–13]. In another experiment [9], the migration of T-lymphocytes was studied on a surface with sinusoidal (wavy) height undulations. The cells were found to move axially in the grooves (minima) of the surface topography, avoiding migration on the ridges (maxima). In [10], the dynamics of several cell types was studied on a 2D sinusoidal surface. Adherent fibroblast cells, dominated by stress-fibers and weakly motile, were found to settle in the concave grooves or adhere aligned to the undulation axis (both on grooves and ridges) [14, 15]. In many adherent cells, the alignment is found to be determined by the competition between the bending energy of the stress fibers, of the nucleus and the contractile forces [16–18]. At the level of cell collectives, both alignment and cell migration within the confluent tissue, is found to be affected by the substrate curvature, experimentally [19–24] and in theoretical analysis [25].

Despite these studies, the underlying mechanisms that determine the response of migrating cells to the substrate curvature are still not well understood, even at the level of single migrating cells. A few theoretical studies addressed the curvature response of an isolated motile cell. One model contains a detailed description of the cellular mechanics, and is based on the assumption of a central role for the nuclear dynamics in controlling the cell migration on the curved surface [26]. A similar approach of modelling cell migration as arising from coupling the nucleus to random peripheral protrusions [27], produced migration patterns that were in qualitative agreement with observations [9], i.e. resulting in cells migrating preferentially along the grooves. Another model provides a simpler and more general description in terms of an active-fluid [28], but its predictions were not systematically compared to experiments. A similar model was proposed to describe amoeba cells moving along ridges, guided by a reaction-diffusion mechanism adapted from macropinocytic cup formation [29].

For cell migration that is driven by the lamellipodia protrusion, understanding the migration on curved substrates requires an understanding of the mechanisms that drive the formation of the lamellipodia. Recently we have proposed a theoretical model where the lamellipodia forms as a self-organization of curved actin nucleators, coupled with adhesion to the substrate [30, 31]. We showed that due to the spontaneous curvature of the actin nucleators, they aggregate at the cell-substrate contact line, and induce an outwards normal force, which represents the protrusive force due to actin polymerization. Curvature sensitive membrane complexes that contain actin nucleation factors [32–35] have been found in experimental observations at the leading edge of cellular protrusions [36–39]. This model can give rise to the spontaneous formation of a lamellipodia-like protrusion, with a stable and asymmetric leading edge, that drives the migration of the simulated membrane vesicle (Fig.1). We found these motile vesicles to be highly persistent on a flat surface, maintaining robustly their direction of migration [31].

**FIG. 1.**
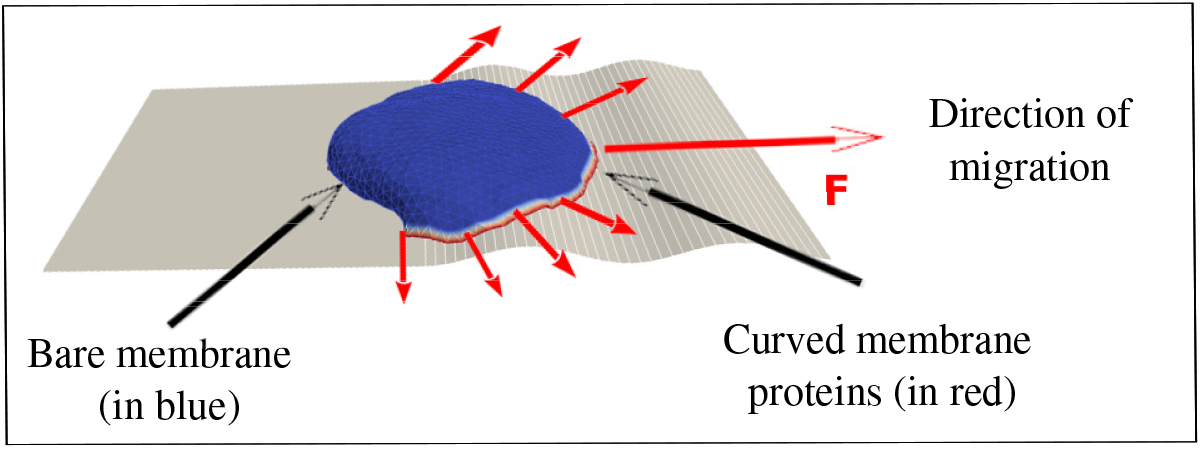
Motile vesicle migrating on sinusoidal substrate. The red dots on the rim of the migrating vesicle are the curved membrane protein complexes with positive intrinsic curvature (convex), while the blue part represents bare membrane. The red arrows with filled arrowheads indicate the forces exerted by the curved protein complexes on the membrane, directed towards the local outwards normal. The total force is indicated by the red arrow with the empty arrowhead, which gives the net propulsion force, and direction of migration of the vesicle.

Here, we use the spontaneously migrating vesicle that arises in our model (Fig.1), as a minimal model of a migrating cell, to explore its behavior and motility on a wide range of curved surfaces. Indeed, we explore surfaces with smooth sinusoidal shape undulations, as well as fibers (outside of cylindrical surfaces) and tubes (inside of cylinders). We do not explore here topographies with sharp edges and barriers or on length-scales much smaller than the cellular length-scale (such as these experimental studies [40–42]), as these will require a much finer mesh for the vesicle surface triangulation and are consequently computationally costly. In addition, sharp edges will increase the chance of our motile vesicle loosing its polarity [31].

Despite the simplicity of our model, the migration patterns of our motile vesicle on the curved surfaces correspond closely to published, as well as new experimental observations that we present here, of cell migration over curved surfaces. The model vesicle is found to move perpendicular to a sinusoidal topography of short length-scale, while it tends to migrate circumferentially around fibers (and pillars). These calculated migration patterns of the motile vesicle, are used to predict the migration patterns of motile cells, and we verify these predictions in experiments using several cell types. Our minimal model for cell migration suggests that some aspects of curvotaxis, of cells that migrate using lamellipodia protrusions, can be universally explained using physical principles.

## II. RESULTS

When simulating the migration of our motile vesicle (Fig.1) on curved surfaces, we have to note that our motile vesicle can easily loose its polarization and motility if it encounters large amplitude and sharp height undulations or barriers [31]. This “fragility” of the motile phenotype in our model constrains us to explore surfaces with small gradients of height undulations, such that our vesicle does not loose its polarization and motility. Our vesicle loses its motility when its leading edge protein cluster (Fig.1) breaks up into two or more parts, which happens when the vesicle collides with an obstacle of large height gradient, or can occur spontaneously due to noise [31]. In our model this event is irreversible, while in real cells there are internal mechanisms that allow cells to recover their polarized shape and resume motility [43–45].

We therefore explore below the migration of our motile minimal-cell system on smooth surfaces where the curvature changes gradually on the length-scale of the vesicle surface triangulation. The shape of the sinusoidal substrate in the simulation is of the form: *z* = *z*_*m*_ sin(2*πy/y*_*m*_), such that the sinusoidal variations are along the y-direction and the curvature remains constant along x-direction. We use several combinations of *z*_*m*_ and *y*_*m*_ in our simulations: (1) *z*_*m*_ = 10 *l*_*min*_; *y*_*m*_ = 120 *l*_*min*_, (2) *z*_*m*_ = 2 *l*_*min*_; *y*_*m*_ = 30 *l*_*min*_, and (3) *z*_*m*_ = 1 *l*_*min*_; *y*_*m*_ = 15 *l*_*min*_. By keeping the ratio *z*_*m*_*/y*_*m*_ *≪* 1, we remain in the regime of small undulations, which maintains the motility of our simulated vesicle.

Similarly for the migration on fibers and inside tubes, we keep their radius large compare with the triangulation length-scale. See Sec. S1 for more details regarding the theoretical model [30, 31, 46–50].

In the simulations with sinusoidal surface, we start with a motile vesicle, that we formed on a flat substrate, and then deform the substrate into a curved shape and allow the vesicle to evolve so that it matches the curved substrate to which it is adhered (see SI Sec. S3, Fig.S3 and Movie-S1 for more details). This allows us to control the initial direction of motion of the vesicle. Alternatively, we can start from a spherical-like vesicle and let it spread over the curved surface. However, in this case, we do not have any control over the initial direction of migration.

### A. Cellular migration on sinusoidal surfaces: large wavelength

We start by studying the vesicle migration on a sinusoidal substrate where the sinusoidal wavelength is larger than the diameter of the adhered cell, and the vesicle can migrate while it is roughly in a region with only one type of substrate curvature over its entire contact surface with the substrate: the groove/ridge width *y*_*m*_*/*2 is about twice larger than the cell diameter 2*R*_*vesicle*_. In Fig.2(i,ii) we show the configurations and trajectories of a motile vesicle that was placed initially either on the bottom (Fig.2A) or top (Fig.2B) of the sinusoidal surface undulation. The vesicle is initially aligned parallel to the surface undulations (along the *x*-axis).

**FIG. 2.**
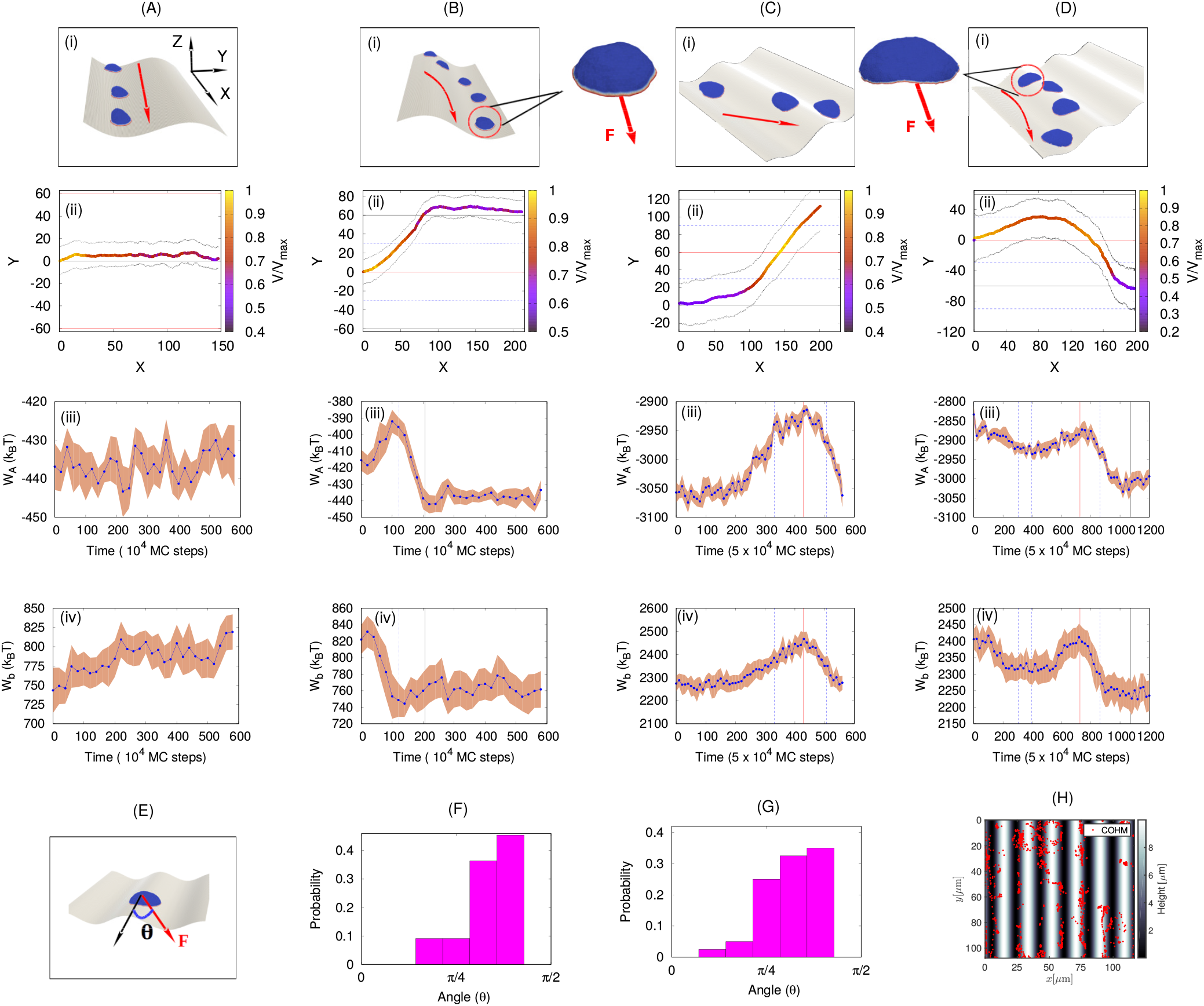
Motile vesicle moving on a sinusoidal substrate with *y*_*m*_*/R*_*vesicle*_ *≫* 1. We use *z*_*m*_ = 10 *l*_*min*_; *y*_*m*_ = 120 *l*_*min*_ for sinusoidal substrate. (A) A small vesicle starting from the minimum of the sinusoidal substrate continues to migrate along its initial direction of migration. (B) Small vesicle starting from the maximum of the sinusoidal substrate shifts to the minimum of the substrate. (C) Large vesicle starting from the minimum of a sinusoidal substrate (with *F* = 2.0*k*_*B*_*T/l*_*min*_), crosses the maximum and reaches the next minimum. (D) Large vesicle starting from the maximum of a sinusoidal substrate (with *F* = 1.0*k*_*B*_*T/l*_*min*_) initially tends to migrate along the positive *Y* -axis, then changes its direction of migration towards the negative *Y* -axis, and finally reaches the minimum. The panel (i) shows the snapshots (with red arrows showing the direction of migration), panel (ii) shows the trajectories, panel (iii) shows the adhesion energy with time and panel (iv) shows the bending energy with time. (E) We define migration angle (*θ*) as the angle between the direction of migration of the vesicle (towards the net active force *F*) and the axis of the sinusoidal substrate (*x*-axis). (F) The distribution of angle at which the vesicle crosses the maxima, generated from simulation. Here we only use the data for large vesicle, as small vesicle in this case does not cross the ridges. (G) The distribution of angle at which the vesicle crosses the maxima, generated from the experimental trajectories of Ref. [9]. (H) The accumulated positions of center-of-mass of *Dictyostelium discoideum* cells over time. For small vesicle (A-B), we use *N* = 607, *E*_*ad*_ = 3.0 *k*_*B*_*T, F* = 4.0*k*_*B*_*T/l*_*min*_ and *ρ* = 4.9%. For large vesicle (C-D), we use *N* = 3127, *E*_*ad*_ = 2.0 and *ρ* = 2.4%.

When we use a vesicle of small size (Fig.2A,B) we find simple dynamics on the sinusoidal surface: when initiated inside the groove, it maintains its aligned direction of motion (Fig.2A, Movie-S2). When initiated on the ridge, the vesicle quickly reorients to almost perpendicular direction of motion, slides to the nearby groove, where it resumes its aligned migration (Fig.2B, Movie-S3).

A larger vesicle (surface area 5 times larger) on the same sinusoidal substrates exhibits more complex dynamics (Fig.2C,D; Movie-S4,S5). This is due to the vesicle now extending over a larger surface and simultaneously spanning more of the two signs of the substrate curvatures. For example, when started in the groove (Fig.2C(i,ii)), it is affected by the nearby ridge, which causes a reorientation similar to that observed in Fig.2B(i,ii). Occasionally, the larger vesicle remain aligned in the groove (see Fig.S4A), but its leading edge aggregate often breaks up, sometimes leading to a loss of the motile phenotype. When the larger vesicle is initiated on the ridges it reorients towards the nearby groove, but due to spanning both sides of the ridge, the vesicle can change its direction during this process (Fig.2D(i,ii)). More examples of these dynamics are shown in SI Sec. S4, Fig.S4 (also see Movie-S6,S7,S8).

In order to understand this behaviour, we plot the adhesion (*W*_*A*_, Eq.S6) and bending energies (*W*_*b*_, Eq.S1) of the vesicle as it is moving between the ridge and groove regions (Fig.2A-D(iii, iv)). We note that both the adhesion and bending energies are roughly constant when the vesicle migrates in the groove. When the vesicle shifts from the ridge to the groove, both the adhesion and bending energies decreases, driving the preference for the vesicle to remain inside the groove. This is easy to understand, as the vesicle can adhere more snugly when “filling” the concave groove, with lower bending energy at the vesicle rim, compared to being more curved on the ridge region. On the other hand, the curved nucleators form stronger bonds between themselves (*W*_*d*_, Eq.S4), and therefore a more robust leading edge cluster, when on the ridge (Fig.S5). However, the changes in this energy term are small compared to the changes in the bending and adhesion energy. These observations explain why energetically it is overall more favorable for the vesicle to reside in the grooves, while the more cohesive leading edge cluster gives rise to faster motility when the vesicle crosses the ridges. Note that the cell-substrate adhesion energy was previously identified as the driving mechanism for the tendency of cells to accumulate in concave grooves and pits [26].

Our theoretical results shown in Fig.2(A-D)(i,ii) are similar to the experimental observations of T lymphocytes migrating on sinusoidal surfaces [9]. In these experiments it was found that cells mostly migrate inside, and aligned with the grooves, while occasionally crossing the ridges rapidly and at large angles. The simulations indicate that the vesicle tends to cross the ridges at large angles (Fig.2F), as observed in experiments [9] (Fig.2G). A similar behavior was observed in migrating *Dictyostelium discoideum* cells on a sinusoidal substrate, as shown in Fig.2H [51]. The positions of the center-of-mass of the cells over time shows that this cell type also tends to stay within the grooves, and avoids the ridges. See sec. S2 for the details of experimental methods.

### B. Cellular migration on sinusoidal surfaces: small wavelength

Next we consider the case where the vesicle radius and the wavelength of the sinusoidal undulations are of the same order, so that a vesicle spans both the ridge and the nearby groove(s). Here, we use sinusoidal variations of two types: *z*_*m*_ = 1; *y*_*m*_ = 15 and *z*_*m*_ = 2; *y*_*m*_ = 30 keeping the ratio of *z*_*m*_*/y*_*m*_ fixed.

We find that when we start with a vesicle that is either orthogonal (Fig.3A, Movie-S9) or parallel (Fig.3B, Movie-S10) to the sinusoidal pattern, the vesicle eventually settles to migrate mostly in the orthogonal direction. However, the speed of the vesicle shows clear oscillatory behaviour (Fig.3A,B(iii)). When the vesicle travels from ridge to groove, it moves towards lower energies and thereby moves faster, while in the opposite case, it slows down. These speeds seem to be periodic along the orthogonal direction to the grooves and ridges of the sinusoidal pattern. In Fig.S6(A-B) we show the average speed of the migrating vesicle at different positions of its center-of-mass between two maxima of the sinusoidal pattern, showing clear periodicity.

Note that when the vesicle was aligned inside the groove (initial condition in Fig.3B), it shows some tendency to persist inside the groove. This tendency gives rise to staircase-like trajectories when the vesicle moves at some oblique angle with respect to the sinusoidal pattern (Fig.3B(ii)). We show more simulations of this type in Fig.S7 (also see Movie-S11,S12,S13,S14).

We compare these simulations to experiments using fish keratocytes migrating on sinusoidal substrates [7], with similar ratio of cell size and sinusoidal wavelength (Movie-S15,S16). Fish keratocytes is a perfect cellular system to be compared to vesicles since they are persistent and polarized cells that contain a large lamellipodium driven by protrusive forces exerted by actin polymerization. In Fig.3C,D, we show two typical trajectories, where the cell migrates in a staircase-like trajectory (Fig.3C(i,ii)) or leaves the groove and moves orthogonal to the pattern (Fig.3D(i,ii); Fig.S8(B-D)). The speed of the cell shows similar oscillatory behaviour as observed in our simulation (Fig. 3C,D(iii)). However, due to the noisy cell speed extracted from the experimental trajectories, we could not identify a clear relation between the mean speed and the cell position within the sinusoidal pattern (Fig.S6(C-D)). The experimental speed can be affected by stick-slip cellular retractions and inhomogeneities in the cell-substrate adhesion, which are absent in the simulations. In Fig.S8 we show more experimental trajectories of migrating keratocytes on the sinusoidal substrate, similar to Fig.3C,D (also see Movie-S17,S18,S19,S20).

**FIG. 3.**
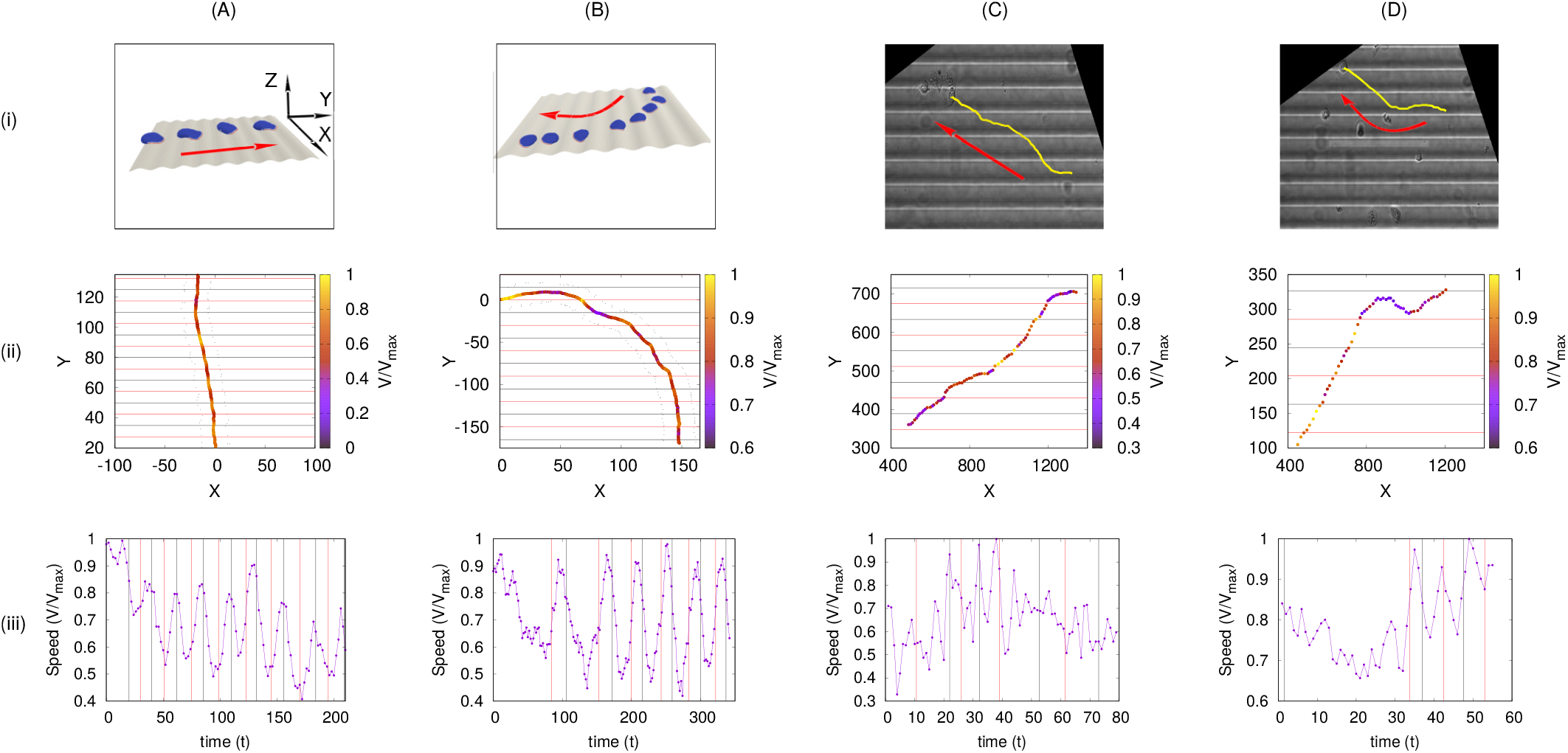
Motile vesicle moving on a sinusoidal substrate with *y*_*m*_*/R*_*vesicle*_ *∼* 1. In this case, we only use small vesicle for simulation results. (A) A vesicle starting in the orthogonal direction of the sinusoidal substrate continues to migrate in the same direction. Here, we use *z*_*m*_ = 1 *l*_*min*_; *y*_*m*_ = 15 *l*_*min*_ for the sinusoidal substrate. (B) A vesicle started from the maximum of the sinusoidal substrate slowly changes its migration direction and becomes orthogonal to the sinusoidal axis. Here, we use *z*_*m*_ = 2 *l*_*min*_; *y*_*m*_ = 30 *l*_*min*_ for the sinusoidal substrate. (C-D) Migrating *karatocytes* on sinusoidal substrate. Here, (i) shows the snapshots (red arrows are showing the direction of migration), (ii) shows the trajectories, and (iii) shows the variation of speed of the vesicle/cell with time. For simulation results, we use *N* = 607, *E*_*ad*_ = 3.0 *k*_*B*_*T, F* = 4.0*k*_*B*_*T/l*_*min*_ and *ρ* = 4.9%.

Despite the favorable comparisons between the model and the experiments on sinusoidal surfaces, it is not easy to interpret the details of the migration process on these surfaces since they contain curvatures of opposite signs. We next explore the migration pattern of our model vesicle, and living cells, on simpler curved surfaces of uniform curvature.

### C. Migration outside and inside a cylindrical surface (fiber and tube)

In order to gain a deeper understanding of the curvature-dependent motility in our model, we simulate the vesicle motion on a surface of uniform curvature, such as the convex curvature of the external surface of a cylinder (fiber). In Fig.4A we plot the dynamics of a motile vesicle, when initially it was aligned along the axis of the fiber (of radius *R* = 10 *l*_*min*_). We find that the vesicle spontaneously shifts its orientation, and ends up rotating along the circumferential direction, as the final steady-state of the system (Fig.4A, Movie-S21). This tendency, to polarize and migrate perpendicular to the axis of the fiber, explains naturally the tendency of the vesicle to migrate perpendicular to the undulation pattern, when moving over the ridges of the sinusoidal surfaces (Figs.2,3).

We can understand the driving force for this re-orientation of the migration, by plotting the adhesion, bending and protein-binding energies of the vesicle during this process (Fig.4B-D) as a function of the migration angle, which is defined as the angle between the direction of motion and the fiber axis (see SI Sec. S9,S10; Fig.S9,S10 for more details). We see that there is a small gain in adhesion (decrease in adhesion energy), decrease in overall bending energy, and a small decrease in the protein binding energy. When oriented circumferentially, the leading edge active forces can stretch the vesicle sideways along the cylinder’s axis, which is efficient in increasing the adhered area along a direction of low curvature, by keeping the membrane close to the fiber surface (Fig.4E). By comparison, when the vesicle is oriented along the axis (Fig.4F) only a small region of the leading edge, along the axis, can pull the membrane close to the fiber and maintain its adhesion. The parts of the leading edge that point along the circumferential direction are less effective in increasing the adhered area due to pulling the membrane off the surface, as well as increasing its bending energy.

This predicted tendency for cells to rotate around fibers, when their migration is driven by a lamellipodial protrusion, is nicely verified by experimental data on *Dictyostelium discoideum* cells [11]. The observed trajectories of migration are biased along the circumferential direction (Fig.4G, Movie-S22), as shown by the peak in the distribution of cellular migration direction (Fig. 4I). The speed of the cells was also found to be maximal along the circumferential direction (Fig. 4H).

**FIG. 4.**
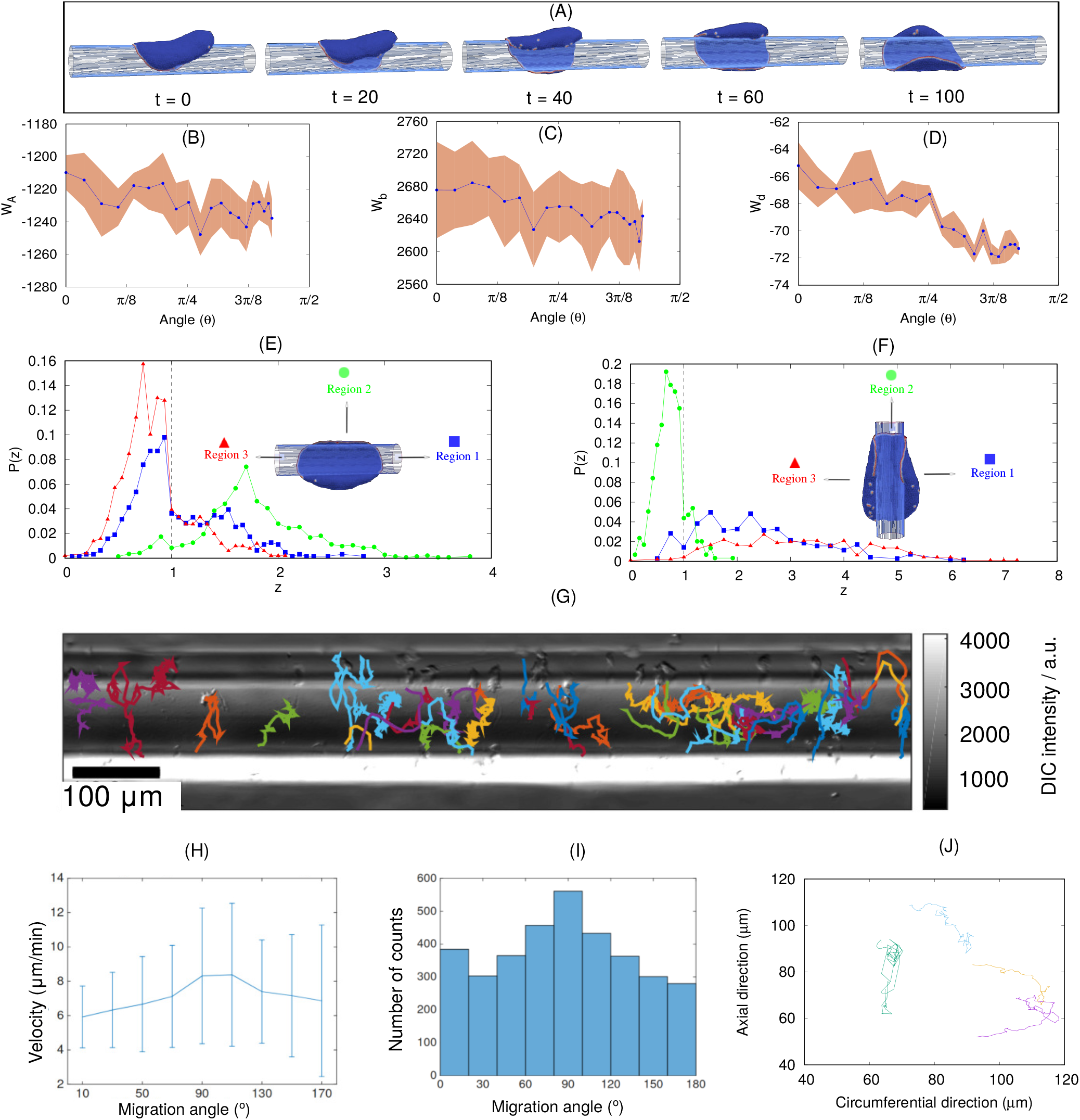
Vesicle migrating outside of a cylindrical fiber. (A) Configurations of the motile vesicle migrating on a fiber, initially in the axial direction, and finally reorients to rotate circumferentially. (B) The adhesion, (C) bending, and (D) binding energies of the vesicle as function of its migration angle during the reorientation process shown in (A). (E) Distribution of the distance *z* of a curved protein along the leading edge from the cylindrical surface, when the vesicle is oriented circumferentially. We plot this distance distribution for three different sections of the leading edge, along three directions, as defined in the inset. The part of the distributions that are on the left side of the vertical dashed line (at *z* = 1) represent adhered proteins. (F) Same as (E), when the vesicle is oriented axially. (G) Trajectories of different D.d. cells on the fiber of diameter 160 *µm* [11]. (H) The distribution of migration speeds of D.d. cells, as function of its migration angle. (I) Distribution of migration angles of D.d. cells on the fiber. (J) The trajectories of MDCK cells migrating on a fibers of 50 *−* 70 *µm* in diameter. For simulation results, we use *R* = 10 *l*_*min*_, *F* = 2.0, *E*_*ad*_ = 1.0 and *ρ* = 2.4%.

Furthermore, it was found experimentally that the tendency of the cells to migrate circumferentially decreased as the fiber radius increased (Fig.S11) [11]. Our model can offer an explanation of this trend, as we find that the energetic advantage of the circumferential orientation in our simulations decreases with increasing fiber radius.

A similar tendency was observed for motile MDCK cells on a fiber (Movie-S23). The trajectories in Fig. 4J show that these cells were either migrating in fast and highly directed bursts along the circumferential direction (Fig.S12), or moving slowly in random motion along the axial direction. This agrees with the model’s prediction that the lamellipodia’s leading edge is more robust along the circumferential direction, which should result in more persistent motion in this direction.

Previous studies with MDCK cells moving on very thin fibers (fiber cross-section circumference same or smaller than the cell diameter) reported a bi-phasic migration pattern [19]. Isolated cells were sometimes observed to migrate axially with high speed, and with a very small adhered surface area. The cell body in these cases exhibits a highly rounded shape, typical of cells under strong contractile forces. Such contractile forces are outside the present model, and we therefore do not expect to reproduce this axially motile phenotype [52]. However, a second phenotype was observed in these experiments, when cells spread and adhere strongly to the fiber surface. During these times the cells seem to exhibit short rotation periods around the fiber circumference [19], but their duration were too short to be conclusive. In addition, the overall orientation of the actin fibers in a confluent monolayer of cells on the fiber was found to be circumferential, in agreement with the orientation of the isolated cell in our simulations.

Another example for spontaneous rotational migration of cells on cylindrical surfaces, is shown in Fig.5(A-C) (Movie-S24,S25). Here *Dictyostelium discoideum* cells are shown to rotate persistently on the external surface of pillars with circular cross-section. On pillars with triangular cross-section, we find that the cells slow down periodically whenever they cross the higher curvature corners (5(D,E)).

**FIG. 5.**
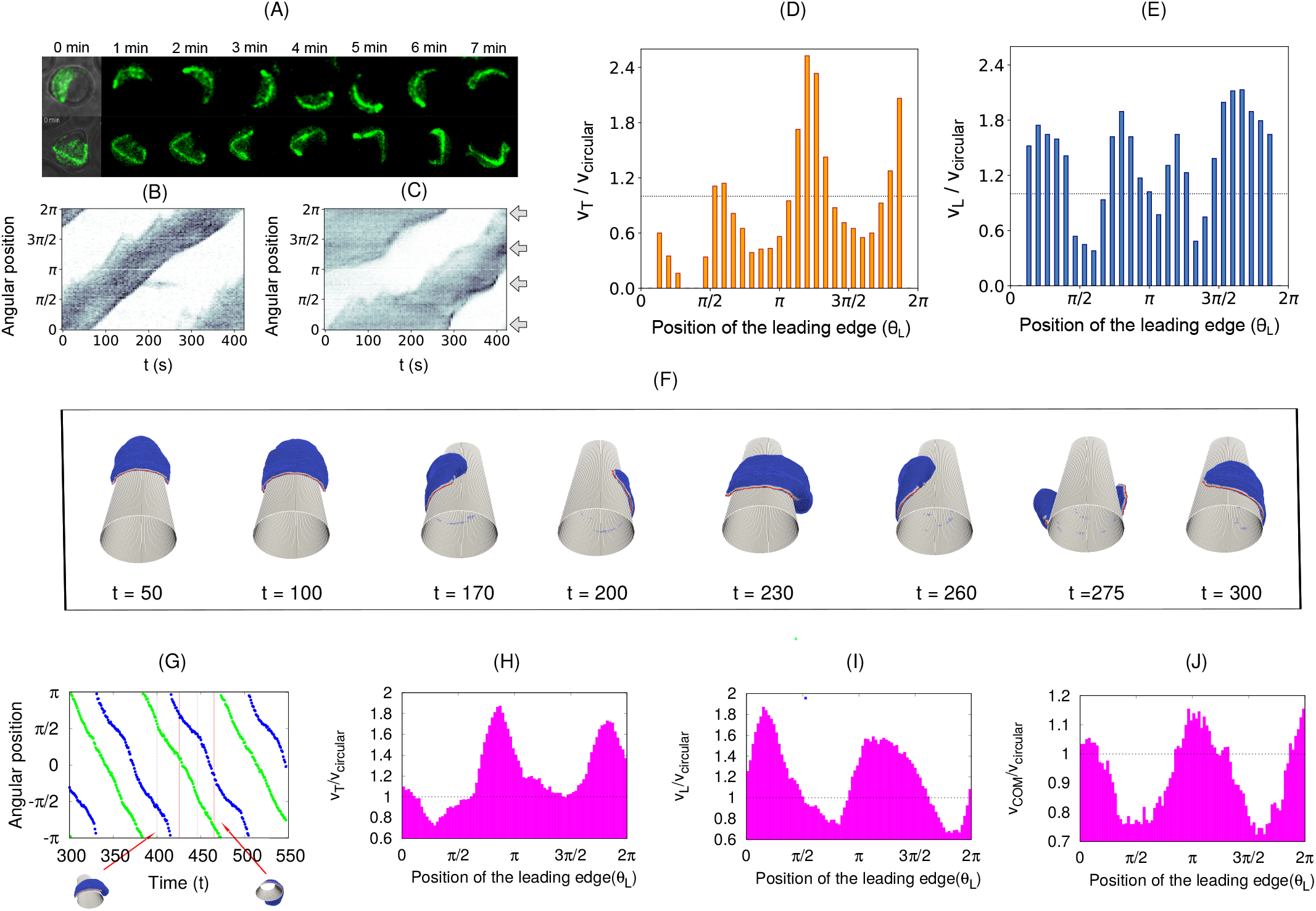
Curvature sensing on micropillars. (A) Timelapse snapshots of *Dictyostelium discoideum* cells moving along the surface of a circular (top) or triangular (bottom) shaped micropillar. In both cases, the field of view shown is 30 × 30 um. Cells express LifeAct-GFP. (B) Kymograph of the actin signal for the cell shown in (A) on the round micropillar. The intensity has been integrated in each angular slice and colour coded from white (0) to black (max). (C) Same as (B) for the triangular shaped pillar. The arrows indicate the positions where the triangle corners are located. (D) Variation in the speed of the trailing edge *V*_*T*_ (scaled by the average speed for the fiber with circular cross-section) as a function of the position of the leading edge (*θ*_*L*_). (E) Variation in the speed of the leading edge (*V*_*L*_) as a function of the position of the leading edge (*θ*_*L*_). (F) Snapshots of the migrating vesicle on a fiber with elliptical cross-section with aspect ratio *r* = 1.54. The vesicle initially migrates in the axial direction and finally reorients along the circumferential direction for longer time (*t >* 200). (G) Kymograph of the leading edge (blue circles) and the trailing edge (green circles) when the vesicle is rotating circumferentially (*t >* 300). (H) Speed of the trailing edge of the vesicle *V*_*T*_ (scaled by the speed for the fiber with circular cross-section *V*_*circular*_) as a function of the position of the leading edge (*θ*_*L*_) over a full period. (I) Speed of the leading edge of the vesicle *V*_*L*_ as a function of the position of the leading edge (*θ*_*L*_) over a full period. (J) Speed of the centre of mass (com) of the vesicle *V*_*COM*_ as a function of the position of the leading edge (*θ*_*L*_) over a full period. For simulation, we use *R*_*x*_ = 12 *l*_*min*_, *R*_*y*_ = 7.773 *l*_*min*_, *E*_*ad*_ = 1.5 *k*_*B*_*T, F* = 2.0 *k*_*B*_*T/l*_*min*_, and *ρ* = 2.4 %.

Note that when cells are migrating on extremely thin fibers, the motility mode is very different, driven by elongated and thin protrusions on either side of the cell [52]. There is no single lamellipodium that drives the migration, and no global rotation of the cell around the fiber. However, the leading edges of the protrusions tend to coil around the fiber. We suggest that this behavior is driven by the same mechanism that we identified here to cause global rotations on larger fibers [47].

In order to verify the above predictions, we simulated the migration of the motile vesicle on a fiber of elliptical cross-section, such that the rotating vesicle experiences different curvatures periodically (Fig.5F, Movie-S26). By plotting the kymograph (Fig.5G) and the speed as function of position (Fig.5H-J), we find periodic variations in the speed of the vesicle that are similar to those observed in the experiments (Fig.5B-E). Note that the experiments exhibit three peaks, due to the triangular shape, compared to two peaks in the simulations on the elliptic cross-section.

In the SI Sec. S13, Fig.S13, we compare the dynamics of migrating vesicles on elliptical fibers of different aspect ratio *r* = *R*_*x*_*/R*_*y*_ (Movie-S27,S28). We note that it takes more time for the vesicle to reorient towards the circumferential direction as the aspect ratio *r* increases (Fig.S13 A,B). For the largest aspect ratio that we tested (*r* = 2.87), the vesicle does not reorient at all (Fig.S13 C), due to the sharp corners that present bending energy barriers. This inhibition of rotation over the sharp corners is similar to the inhibition of coiling at the leading edge of cellular protrusions, calculated and observed when cells spread over fibers [47].

Finally, we simulated the migration of the motile vesicle inside a cylindrical tube (see SI sec. S14, Fig.S14,S15, Movie-S29,S30). We find that the vesicle prefers to migrate along the tube axis, but easily loses its motility, and can even form “bridges” across the tube axis. This prediction was verified experimentally for MDCK cells, which were found to be weakly motile, with the most persistent motility periods aligned with the tube axis or spread across the tube axis (Fig.S15G, Movie-S31).

## Supporting information

Supplementary informations

## III. CONCLUSION

We demonstrated here that a minimal physical model of a motile cell, based on very few ingredients and energy terms, is able to describe and explain several curvotaxis features of lammelipodia-based cell migration on curved adhesive substrates. Within this minimal model, cell migration arises when a leading edge cluster of highly curved membrane protein complexes forms, due to these protein complexes being both highly curved, binding to each other and exerting protrusive forces on the membrane. These forces represent in the model the pressure exerted on the membrane when actin polymerization is initiated at the membrane by these curved protein complexes, which contain actin nucleation factors (such as WAVE [53–55]). The curvotaxis features that the model explains, such as the tendency of motile cells to migrate aligned within grooves, avoid ridges, and rotate around fibers, all arise due to minimization of the adhesion and bending energies of the vesicle. The advantage of simple, physical models is demonstrated here, exposing general mechanisms that are universal and not cell-type-specific.

The curvotaxis property of the motile “minimal-cell” is shown to be a truly emergent phenomenon of the whole motile vesicle. Within our model, the energy minimization that aligns the migration of the “minimal-cell” arises from shape changes of the whole vesicle in response to the imposed curved surface and the organization of the curved membrane complexes that form the leading-edge cluster (and motility). The curved membrane complexes are sensitive to curvature on a much smaller length-scale compared to the cell-size, and therefore do not directly determine the preferred curvotaxis response of the whole motile vesicle.

Eukariotic cells contain numerous additional components that our simple model does not contain, such as the effects of contractility, stress-fibers and internal organelles (such as the big nucleus), which can all affect migration on curved substrates. Nevertheless, the agreement between the predictions of the model and the observations of curvotaxis in different types of motile cells, suggests that these simple energetic considerations may drive curvotactic features in cells, despite the biochemical complexity and differences between cells. These results demonstrate that complex cellular behavior may have physical underpinnings, with added layers of biological complexity and regulation. The framework presented here could serve in the future to explore cell migration in more complex geometries [10, 56], and over soft substrates (such as other cells) with dynamic curvature.

## IV. AUTHOR CONTRIBUTIONS

R.K.S., S.P., A.I. and N.S.G. developed the theoretical model; R.K.S. and N.S.G. conceived, designed and implemented the analysis of the model, and prepared the manuscript; M.L. and S.G. conceived and performed the experiments on keratocytes migrating on sinusoidal substrate and contributed the data; W.X. and B.L. performed the experiments and analysis of the MDCK cells migrating inside the tube and outside of fiber and contributed the data; M.S., Christoph Blum, M.T., O.S. and E.B. conceived and supervised the experiments on D.d. cells migrating on fiber and sinusoidal substrate and contributed the data; Carsten Beta and C.M.T. conceived and supervised the experiments on D.d. cells migrating on pillars and contributed the data; All authors reviewed and edited the manuscript.

## V. ACKNOWLEDGMENT

We thank Junsang Doh and collaborators for providing the data for the migration of T lymphocytes of sinusoidal wavy surfaces. N.S.G. is the incumbent of the Lee and William Abramowitz Professorial Chair of Biophysics, and acknowledges support by the Ben May Center for Theory and Computation, and the Israel Science Foundation (Grant No. 207/22). This research is made possible in part by the historic generosity of the Harold Perlman Family. R.K.S. acknowledges the support from ANR (ANR-19-CE11-0002-03). A.I. and S.P. were supported by the Slovenian Research Agency (ARRS) through the Grant No. J3-3066 and J2-4447 and Programme No. P2-0232. This work was supported by the Marie Sk-lodowska-Curie Actions, Individual Fellowship, Project: 846449 (to W.X.) and the “Initiatives d’excellence” (Idex ANR-11-IDEX-0005-02) transverse project BioMechanOE (TP5) (to W.X.). This work was supported by the European Research Council (Grant No. Adv-101019835 to B.L.), LABEX Who Am I? (ANR-11-LABX-0071 to B.L. and W.X.) and the Ligue Contre le Cancer (Equipe labellisée 2019 to B.L. and W.X.) and the ANR PRC LUCELL grant (ANR-19-CE13-0014-01 to B.L. and W.X.), and DIM-ELICIT 2019: Equipment support, Région Ile-de-France (to W.X. and B.L.). B.L. and W.X. acknowledge the ImagoSeine core facility of the IJM, member of IBiSA and France-BioImaging (ANR-10-INBS-04) infrastructures. The research of Carsten Beta and C.M.T. has been partially funded by the Deutsche Forschungsgemeinschaft (DFG), Project-ID No. 318763901—SFB1294. S.G. acknowledges funding from FEDER Prostem Research Project no. 1510614 (Wallonia DG06), the F.R.S.-FNRS Epiforce Project no. T.0092.21, the F.R.S.-FNRS Cellsqueezer Project no. J.0061.23, the F.R.S.-FNRS Optopattern Project no. U.NO26.22 and the Interreg MAT(T)ISSE project, which is financially supported by Interreg France-Wallonie-Vlaanderen (Fonds Européen de Développement Régional, FEDER-ERDF). M.L. is financially supported by the WBI-World Excellence Grant Programme for long-term scholarship.

## Notes

### Competing Interest Statement

The authors have declared no competing interest.

