## Supplementary informations for "A minimal physical model for curvotaxis driven by curved protein complexes at the cell’s leading edge"

**This PDF file includes:**

Supplementary Text

Secs. S1 to S14

Figs. S1 to S16

Legends for Movies S1 to S31

References (1-30)

**Other Supplementary Materials for this manuscript include the following:**

Movies S1 to S31

#### S-1. THEORETICAL MODEL

The migrating cell is represented in our theoretical model using a three-dimensional membrane vesicle. The vesicle is described by a closed triangulated surface having $N$ vertices, connected to their neighbours with bonds, and forming a dynamically triangulated, self-avoiding network, with the topology of a sphere [1–7] (Fig.S-1). The nodes that compose the vesicle surface can either represent the bare membrane (blue in Fig.S-1), or represent membrane protein complexes with convex spontaneous curvature [8], that diffuse on the membrane surface, having nearest-neighbor attractive interaction with each other (red in Fig.S-1). Convex protein or membrane curvature stands for a node that is locally protruding outwards, with respect to the vesicle interior.

We consider that each curved protein complex recruits actin polymerization, which gives rise to a local protrusive force that pushed the membrane. This is represented in our model as an active force ($F$) exerted at the site of the curved protein on the membrane, in the direction of the local outward normal to the vesicle surface. The simplifying assumption is that the actin polymerization that occurs near the membrane can be treated as a local force exerted directly at the site of the curved protein complex which includes actin nucleation factors (such as the WAVE complex [9–11]).

The vesicle energy has therefore the following contributions: The continuum version of the bending energy, is given by,

[a]
[b] Present address: Institut Curie, PSL Research University, CNRS, UMR 168, Paris, France
[c]

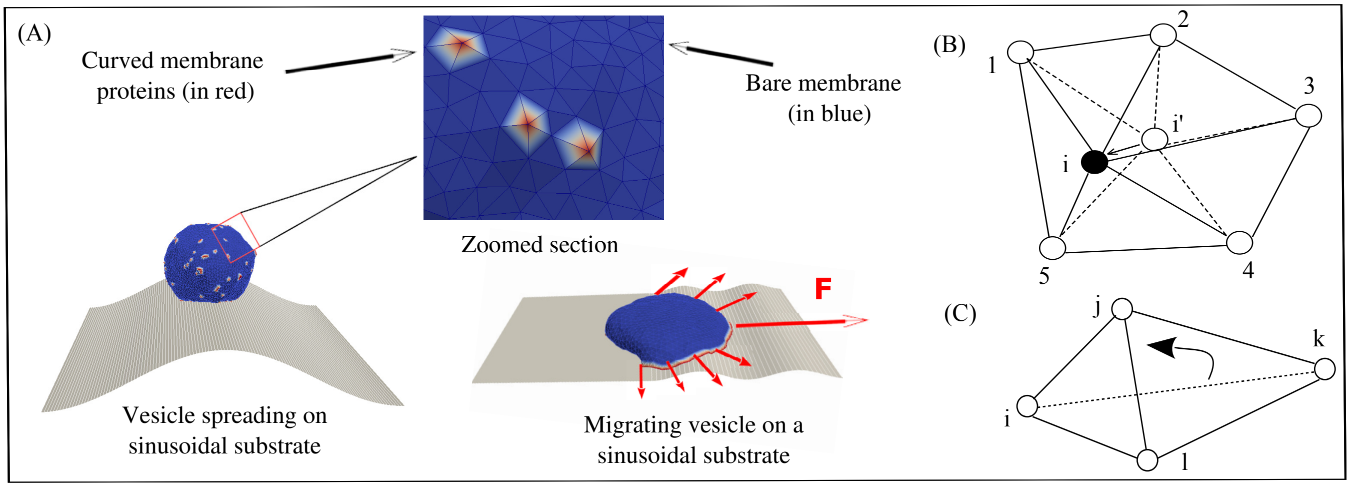


FIG. S-1. Schematic representation of our model. (A) The vesicle is formed by a closed triangulated surface, having $N$ vertices connected to its neighbours with bonds. These bonds can change their length such that they are never below $l_{min}$, or above $l_{max} = 1.7l_{min}$. The red dots on the surface of the vesicle represents the curved membrane protein complexes with positive intrinsic curvature (convex), while the blue part represents bare membrane. A zoomed version of a small section of the vesicle surface is shown in the inset. We show here two possible initial conditions: (left) The vesicle starts with a spherical-like shape, adheres, spreads and migrates on the curved surfaces. (right) We generate a motile (crescent shaped) vesicle on a flat substrate and then deform the surface and let the vesicle evolve to conform to the deformed (curved) shape, and then allow it to migrate. (B) Vertex movement: The vertex $i'$ is moved to $i$. (C) Bond flip: The bond $i-k$ is flipped to bond $j-l$.

$$W_b = \frac{\kappa}{2}\int_A (H - H_0)^2 dA, \tag{S-1}$$

where $\kappa$ is the bending rigidity, $H$ is the mean local curvature of the membrane surface, $H_0$ is the local spontaneous curvature, and the integral is over the entire surface. The vesicle contains curvature sensitive protein complexes, that occupy vertices with an overall density $\rho = N_c/N$, where $N_c$ is the number of such protein nodes, and $N$ is the total number of vertices on the vesicle. These protein nodes have a positive (convex) spontaneous curvature ($H_0 > 0$), while the bare membrane nodes have zero spontaneous curvature. In our simulations we use a discrete version of the bending energy [12, 13] which we calculate in the following way:

In the absence of any spontaneous curvature, the integration of the square of the mean curvature over the entire surface can be discretized in the following way [13]:

$$\int_A H^2 dA = \sum_i \frac{1}{\sigma_i}\left[\sum_{j(i)} \frac{\sigma_{ij}}{d_{ij}}(\mathbf{R}_i - \mathbf{R}_j)\right]^2$$

where the outer sum runs over all the vertices and the inner sum runs over all the neighbours of $i$-th vertex. $\mathbf{R}_i$ is the radial vector of the vertex $i$, $d_{ij}$ is the distance between the vertices $i$ and $j$, $\sigma_i$ is the area of the cell (formed by the vertex $i$ and all its neighbours) in the dual lattice defined as,

$$\sigma_i = \frac{1}{4}\sum_{j(i)} \sigma_{ij} d_{ij}$$

where $\sigma_{ij}$ is the distance between the vertices $i$ and $j$ in the dual lattice.

In the presence of curved membrane proteins, the spontaneous curvature of the $i$-the vertex is $c_i$ (say). Then the bending energy of the $i$-th vertex can be written as,

$$W_b(i) = \frac{\kappa}{2}\sigma_i\left(\frac{h_i}{\sigma_i} - c_i\right)^2 \tag{S-2}$$

where,

$$h_i^2 = \left[ \sum_{j(i)} \frac{\sigma_{ij}}{d_{ij}} (\mathbf{R}_i - \mathbf{R}_j) \right]^2$$

Thus, the total bending energy of the vesicle can be written as,

$$W_b = \sum_i W_b(i) \tag{S-3}$$

where, the sum runs over all the vertices of the vesicle.

The direct binding energy between the protein complexes on nearest-neighbour nodes is given by,

$$W_d = -w \sum_{i<j} \mathcal{H}(r_0 - r_{ij}), \tag{S-4}$$

where, $\mathcal{H}$ is the Heaviside step function, $r_{ij} = |\overrightarrow{r}_j - \overrightarrow{r}_i|$ is the distance between proteins, $\overrightarrow{r}_i$, $\overrightarrow{r}_j$ are the position vectors for $i, j-th$ proteins, and $r_0$ is the range of attraction, $w$ is the strength of attraction. The range of attraction is chosen such that only the proteins that are in neighbouring vertices can bind to each other.

These curved protein complexes also recruit actin filaments that polymerize at the location of these proteins. We assume that the direction of these forces are normally outward of the local surface containing the proteins. The active energy is given by,

$$\Delta W_F = -F \; \hat{n}_i \cdot \overrightarrow{\Delta r}_i, \tag{S-5}$$

where, $F$ is the magnitude of the active force, representing the protrusive force due to actin polymerization [2, 5] that is acting in the direction of outward normal vector of the local membrane surface (along $\hat{n}_i$) and $\overrightarrow{\Delta r_i}$ is the displacement vector of the protein complex. The "active" forces in our simulations are implemented as external forces that act on the specific nodes of the system that contain the curved membrane proteins (with positive spontaneous curvature, red nodes in Fig.S-1). This is done by giving a negative energy contribution when the points on which these forces act move in the direction of the force. These forces are "active" since they give an effective energy (work) term that is unbounded from below and thereby drive the system out-of-equilibrium. By exerting a force directed at the outwards normal we naturally describe Arp2/3-driven branching polymerization of actin, which is rather isotropic and acts as a local pressure on the membrane.

Finally, the adhesion energy due to the interaction between the vesicle and the extracellular substrate, is given by,

$$W_A = -\sum_{i'} E_{ad}, \tag{S-6}$$

where $E_{ad}$ is the adhesion energy per node, and the sum runs over all the vertices that are adhered to the substrate [2, 3, 14]. By 'adhered vertices', we mean all such vertices, whose perpendicular distance from the adhesive surface is less than a threshold, which we chose to be equal to the length $l_{min}$, which is the unit of length in our model, and defines a minimal length allowed for a bond. Thus, the total energy of the system is given by,

$$W = W_b + W_d + W_F + W_A \tag{S-7}$$

We update the vesicle with mainly two moves, (1) vertex displacement and (2) bond flip. In a vertex displacement, a vertex is randomly chosen and moved by a random length and direction, with the maximum possible distance restricted by 0.15 $l_{min}$ (Fig. S-1(B)). This movement provides shape fluctuations to the vesicle. In the bond flip move, a single bond is chosen, which is a common side of two neighbouring triangles, and this bond is cut and reestablished between the other two unconnected vertices (Fig. S-1(C)) [2, 3, 14]. The bond flip is responsible for the lateral fluidity of the system that allows the vertices to diffuse through the membrane surface. Since our protein complexes are attached to a particular vertex, it also diffuses along with the vertex in the bond flip movement. The maximum bond length is restricted to $l_{max} = 1.7 \; l_{min}$ in order to maintain self avoidance of triangulated network. We update the system using the Metropolis algorithm, where any movement that increases the energy of the system (Eq.S-7) by an amount $\Delta W$ occurs with rate $\exp(-\Delta W/k_B T)$, otherwise it occurs with rate unity.

### S-2. EXPERIMENTAL METHODS

#### A. Migrating keratocytes on sinusoidal substrate

##### 1. Cell culture

Fish epithelial keratocytes were obtained from the scales of Central American cichlid (Hypsophrys Nicaraguensis) [15, 16]. Scales were gently taken off the fish and placed in the center of a microprinted PDMS-coated glass coverslips and covered with a drop of 150 µL of culture medium. The culture medium was composed of Leibovitz's L-15 medium (Thermo Fisher Scientific) supplemented with 10% fetal bovine serum (FBS, Capricorn), 1% penicillin/streptomycin (Westburg), 14.2 mM HEPES (Sigma Aldrich) and 30% deionized water were put on top of the scale. A glass coverslip of 22 mm in diameter was deposited on top of the scales and few drops of culture medium were added around the samples. Epithelial keratocytes were cultured in the dark at room temperature for 12 h. Keratocytes were detached from the glass coverslip by incubating with a trypsin solution (1 ml per glass slide) for 5 minutes and resuspended in 4 ml of L-15 Leibovitz complete medium. Suspended cells were then transferred to FN-coated corrugated hydrogels. All experiments were made between 2 and 8 hours after cell seeding.

##### 2. Fabrication of corrugated polyacrylamide hydrogels by UV-photocrosslinking

Instead of the standard radical polymerization using catalysts such as tetramethylenediamine (TEMED) and ammonium persulfate (APS), which lead to slow polymerization times, we used an Irgacure 2959 photoinitiator (2-Hydroxy-4'-(2-hydroxyethoxy)-2-methylpropiophenone) to polymerize hydroxypolyacrylamide (hydroxy-PAAm) hydrogels. Hydroxy-polyacrylamide (hydroxy-PAAm) hydrogels were prepared by mixing acrylamide (AAm), bis-acrylamide (bis-AAm), N-hydroxyethylacrylamide (HEA), 2-Hydroxy-4'-(2-hydroxyethoxy)-2-methylpropiophenone (Irgacure 2959, Sigma #410896) and deionized water. A solution composed of 2836 $\mu L$ of acrylamide (AAm, Sigma #79-06-1) at 15% $w/w$ in deionized water, 1943 $\mu l$ of N,N'- methylenebisacrylamide (BisAAm, Sigma #110-26-9) at 2% $w/w$ in deionized water, and 1065 $\mu L$ N-hydroxyethylacrylamide monomers at 65 $mg/mL$ in deionized water (HEA, Sigma #924-42-5) were mixed together in a 15 $mL$ Eppendorf tube [16–18] and deionized water was added to reach a final volume of 6 $mL$. We prepared a stock solution of Irgacure 2959 in sterile deionized water at 5 $mg/mL$. We introduced 1 $mL$ of the stock solution into the 6 $mL$ of the hydrogel solution to obtain a final concentration of 0.7 $mg/mL$. After a gentle mixing, the solution was degassed during 30 $min$ under a nitrogen flow. Glass coverslips of 22 $mm^2$ in diameter were cleaned with 0.1 $M$ NaOH solution during 5 $min$ and then rinsed abundantly with deionized water during 20 $min$ under agitation. Cleaned glass coverslips were then treated during one hour with 3-(trimethoxysilyl)propyl acrylate (Sigma #2530-85-0) to promote a strong adhesion between the hydroxy-PAAm hydrogel and the glass coverslips and finally dried under a nitrogen flow. A volume of 40 $\mu L$ of the degassed mixture was squeezed between an activated glass coverslip and a chromium optical photomask (Toppan photomask, France) and before exposition to UV illumination at 360 $nm$ (Dymax UV light curing lamp). Chromium optical photomasks with alternating transparent stripes of 10 $\mu m$ wide and black stripes of 10 $\mu m$ or 20 $\mu m$ wide were used to form corrugated hydrogels with wavelengths of 20 $\mu m$ ($\lambda$20), 30 $\mu m$ ($\lambda$30) and 50 $\mu m$ ($\lambda$50) and respectively [19]. After UV exposition at 360 nm during 10 min at 10 $mW/cm^2$ through the optical photomask, the polymerization was completed and a corrugated hydroxy-PAAm hydrogel was formed. The amplitude of 20 $\mu m$ ($\lambda$20) and 30 $\mu m$ ($\lambda$30) corrugated hydrogels was changed by adjusting the volume of the degassed polyacrylamide solution squeezed between the glass coverslip and the chromium optical photomask. Finally, hydrogels were gently removed from the photomask under water immersion, washed three times in sterile deionized water under gentle agitation and stored in sterile deionized water at $4°C$. Photocrosslinked hydroxy-PAAm hydrogels were optically transparent and did not exhibit any autofluorescence background at $470 \pm 20$ $nm$, $562 \pm 88$ $nm$ and $591 \pm 21$ $nm$.

##### 3. Time-lapse imaging

Time-lapse microscopy experiments were carried out on a Nikon Ti-U inverted microscope (Nikon, Japan) equipped with Differential Interference Contrast (DIC) mode. Images were taken every 3 min using a ×10, ×20 or ×40 objective and captured with a DS-Qi2 camera (Nikon, Japan) controlled with the NIS Elements Advanced Research 4.0 software (Nikon). Tracking of the indivudal keratocytes on the corrugated hydrogels were performed with CellTracker in semi-automatic mode [20].

##### B. Migration of *Dictyostelium discoideum* (D. d.) cells on sinusoidal and cylindrical substrates

###### 1. Cell culture

Dictyostelium Discoedium (D.D). exists as several cell strains that need to be treated differently. An important difference between strains is the ability to use different food supplies. The cell lines used here were all axenic, meaning that they were able to feed on the culture medium (HL5 (Formedium, Norwich, England)). To use the cells for experiments, frozen stock was thawed at room temperature and afterwards cultured in HL5 medium on Petri dishes. The doubling time of the cells was between 8 to 9 hours at the optimal growing temperature of $21-23°C$. The cell culture was subcultured every $2-3$ days, when the cells have became confluent on in the Petri dish. The passage number was increased by one each time for a new subculture and the cells were discarded after passage 15. In the experiments we used cells harvested in their exponential growth phase. The preparation of the cells started one day before the experiment. 106 cells were pipetted into a flask with 25 $ml$ HL5 medium. This flask is cultivated on a shaking table at $22°C$ with 150 rotation per minute. On the day of the experiment, 7 hours prior to the start of the experiment, the cells were centrifuged and the medium was removed. The cells are washed with phosphate buffer and afterwards centrifuged again. The remaining pellet was diluted with 20 $ml$ phosphate buffer and was positioned on the shaking table at $22°C$. Every 6 minutes a pulse of cAMP (18 Mol; Sigma-Aldrich) was delivered into the shaking culture. After six hours of starvation and the cells were chemotactically competent. AX3-ACA-Null cells were used, as the lack the aggregation stage adenylyl cyclase (ACA), i.e the cells were not able to communicate with each other. This missing functionality enabled us to investigate the effect of the complex geometry without the influence of chemotaxis.

###### 2. Experiments on glass capillaries

The cell migration was observed on glass capillaries. To exclude any communication between the cells by signaling molecules we placed the optical fibers in a perfusion chamber (RC-27, Large Bath Chamber, Warner Instruments, Hamden, CT, USA) on a glass-spacers to allow a fluid flow around the fiber or used a microfluidic device with through flow. Fig. S-2 shows the fiber setup. The cells were imaged from below with an inverted optical microscope through the number 1 cover slip. We use a peristaltic pump (RP-1 Peristaltic Pump, Mettler Toledo Inc., Columbus, Ohio USA) to create a fluid flow with a mean flow speed in the chamber of $v = 167\mu m/s$ . To investigate the actual velocities of the fluid flow close to the fiber, we used fluorescently labeled polymer beads Duke $36-6$ (Polystyrene Divinylbenzene (PS-DVB), Duke Scientific Corporation, California, USA), with a diameter of $33\mu m$. The mean bead velocity close to the fiber was $v = 10\ \mu m/s$. Hence we can be sure that the velocity in the setup did not induce shear driven migration of the D. d. cells but was still high enough to flush away any signaling molecules. The drawbacks of this setup are the lensing effect of the fiber. See Ref. [21] for details.

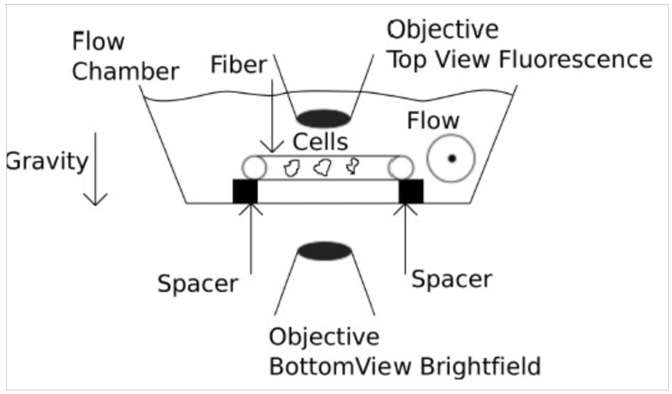


FIG. S-2. Sketch of the curvotaxis setup. The optical fiber is placed on two glass spacers inside the perfusion chamber. The perfusion chamber is connected to a perfusion pump. Cells are added to the fiber from the top. Imaging can be done as well with inverted as with top-view optical setups

##### 3. Experiment on sinusoidal substrate

We use the Photonic Professional (GT) (Nanoscribe) to produce masks for sinusoidal substrates. We chose IP-S, a highly viscous photoresist, in combination with a 25× objective (ZEISS 25×/ 0,8 DIC Imm Korr LCI Plan-NEOFLUAR) and an ITO-coated DiLL glass substrate (size 25 $mm \times 25$ $mm$; thickness 0.7 $mm$; optical transparent; provided by NanoScribe) for the mask production. We use the 3D print as a negative and pour polydimethylsiloxane (PDMS) onto the structure where PDMS and the cross-linker (Sylgard 184, Th. Geyer) are mixed in a mass ratio $10:1$. Finally, the PDMS, as well as an Ibidi bottom glass substrate (ibidi GmbH), are treated with a plasma cleaner provided by Harrick Plasma to clean and to oxidize the surfaces. In doing so, the PDMS sticks well to the substrate after treatment.

The optical setup and D.d. cells are pipetted onto the sinusoidal wave structures. With the help of the spinning disc confocal laser scanning microscope (sdCLSM), the cells are tracked in 3D over time to observe their amoeboid motion. The exposure time is set to $75-100$ $ms$, the acquisition time to $351-376$ $ms$, the z-step is $0.8-1.5$ $\mu m$ and the number of slices is $15-25$. Stacks are recorded every 30 $s$. For details see Ref. [22].

#### C. Spreading and migration of *Madin-Darby canine kidney* (MDCK) cells on fibers and inside tube

##### 1. Microfabrication of elastomeric microtubes and microfibers

Microtubes were fabricated inside polydimethylsiloxane (PDMS) blocks using previously described method [23]. Briefly, a fresh mixture of silicone elastomer base and silicone elastomer curing agent (Sylgard 184, DOWSIL$^{TM}$, $10:1$ by weight) was cured on aligned smooth copper or platinum wires (Goodfellow SARL) of different diameters. The metal wires were later pulled out leaving parallel microtubes in the PDMS block. As-fabricated PDMS blocks were then stuck to a glass-bottom petridish (FluorodishTM, Cat#: $FD35-100$) and coated with fibronectin (Sigma-Aldrich) for cell adhesion.

As described previously [24], PDMS cylindrical microfibers were fabricated by pulling PDMS fibers out of a pre-cured mixture of silicone elastomer base and silicone elastomer curing agent (Sylgard 184, DOWSIL$^{TM}$, $10:1$ by weight). The mixture was mixed and left at room temperature for about 10 h before its viscosity increased to allow pulling fibers. As-fabricated microfibers were hanged in an $80°C$ oven for 1 $h$ for full polymerization. The microfibers were then hanged in a glass-bottom petridish and coated with fibronectin for cell seeding.

##### 2. MDCK cell migration on cylindrical microfibers and microtubes

MDCK-LifeAct-GFP (stable cell line transfected with LifeAct GFP, binding to actin filaments) cells were cultured in complete DMEM medium (Life Technologies), supplemented with 10% fetal bovine serum and 1% penicillin/streptomycin). Cells were cultured at $37°C$ and 5% $CO_2$ conditions until confluent. The cells were then collected and seeded on microfibers and microtubes at 50 million cells/mL. After 2 $h$, the samples were washed carefully with clean DMEM medium to remove un-attached cells and left single, isolated cells attached on the scaffolds.

The samples were then mounted on a confocal microscope (Zeiss, LSM 780). To record a 3D, live-cell video, z-stacks (1 $\mu m$ per Z step) covering the whole volume of PDMS microfibers or microtubes were recorded at 10 min/frame with either 25×, 40× or 63× objectives. 3D time-lapse videos were recorded over a period ranging from 10 to 24 $h$.

For image analysis, we first converted 3D z-stack images into 2D projections as described previously [23]. The 2D time-lapse projections were then used for PIV mapping with PIVlab (an implemented tool for MATLAB R2020) and cell tracking. To avoid bias, we flipped the 2D projected movies so that at the end of the movies, the cells' horizontal positions are on the right side in comparison to their initial position.

#### D. Migration of *Dictyostelium discoideum* (D. d.) cells on micropillars

##### 1. Cell culture and imaging

The non-axenic *D. discoideum* strain DdB NF1 KO [25], transformed with an episomal plasmid encoding for Lifeact-GFP and PHcrac-RFP (SF108, as described in [26]) was used. Cells were cultivated in 10 $cm$ dishes with Sørensen's buffer (14.7 mM $KH_2PO_4$, 2mM $Na_2HPO_4$, pH 6.0) supplemented with 50 $\mu M$ $MgCl_2$, 50 $\mu M$ $CaCl_2$ and using G418 (5 $\mu g/ml$) and hygromycin ($33 \mu g/ml$) as selection markers. Klebsiella aerogenes with an $OD_{600}$ of 20 were added to the solution in 1 : 10 volume to a final $OD_{600}$ of 2. Before imaging, bacteria were removed by washing the cells 3 times with Sørensen's buffer by centrifugation at $300 \times g$. Cells were harvested in the last washing step and transferred to the PDMS block containing the pillar structures. A glass coverslip (#1.5, $24 \times 24$ $mm$, Menzel Glaser) was used to cover the sample and prevent contamination and evaporation of the cell solution. The cells were diluted to a density that enabled imaging of single cells on the surface of the pillars. Temporal recordings were acquired at a rate of 0.2 fps using a laser scanning microscope (LSM780, Zeiss, Jena) with a 488 nm Argon laser and a 40× water immersion objective.

##### *2. Microfabrication of pillar structures*

A silicon wafer was coated with a 10 $\mu m$ photoresist layer (SU-8 2010, Micro Resist Technology GmbH, Germany) and patterned by direct write lithography using a maskless aligner ($\mu$MLA, Heidelberg Instruments Mikrotechnik GmbH, Germany). Polydimethylsiloxane (PDMS, Sylgard 184, Dow Corning GmbH, Germany) at a ratio of 10 : 1 (base to curing agent) was spin coated on the microstructured wafer to obtain a thin ($\sim 300\mu m$) film and cured for 2 $h$ at $75°C$. A PDMS block containing the micropillars design was cut out and placed on top of a glass coverslip (#1, $24 \times 40$ $mm$, Menzel Glaser).

#### S-3. PREPARATION OF MOTILE VESICLES ON CURVED SURFACES

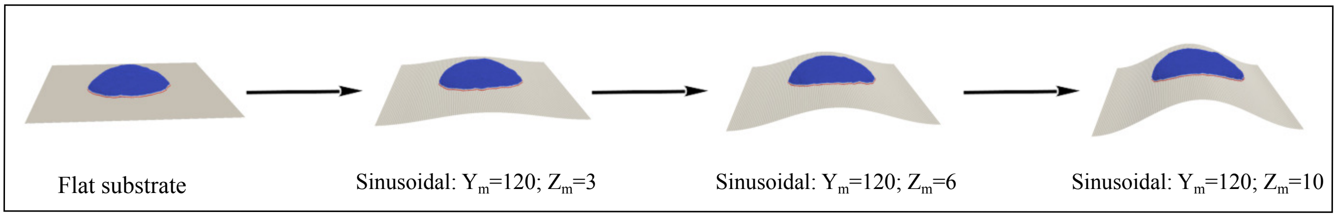


FIG. S-3. Mechanism of preparation of vesicles on curved geometries.

In our simulation, we often obtain a migrating vesicle on curved surfaces (cylinder or sinusoidal) by spontaneous polarization of proteins when started from a spherical-like vesicle. In this case, however, the direction of polarization is random and set by the minimum of energy. So, we could not allow the vesicle to migrate in a direction of our choice. In order to start with a vesicle that migrate in a direction of our choice, we thus use a migrating vesicle already generated on a flat substrate, and place it on a curved surface and allow it to adjust on the curved substrate.

The above mentioned method enable us to study the migration of vesicle of a direction of our choice, but often, the cluster of the migrating vesicle breaks into parts, since at the time of replacing the substrate (flat into curved) some of the vertex containing proteins detaches from the substrate. In order to resolve this issue, we use another method, where we slowly change the curvature of the substrate and allow the vesicle to adjust on the substrate without detaching the vertices from the substrate (Fig. S-3). The process of adhering on the substrate is much faster than the reorientation of the vesicle, such that the vesicle does not rotate much while adhering to the curved substrate.

#### S-4. MORE TRAJECTORIES FOR THE MIGRATION ON SINUSOIDAL SUBSTRATE WITH LARGE WAVELENGTH

Here, we show few more trajectories of vesicle migrating on sinusoidal substrate with large wavelength. In Fig. S-4(A), we show a large vesicle starting from the minimum of the sinusoidal substrate maintains its direction of motion, and finally looses its motility property after long time. Similar behaviour is observed for a small vesicle starting from the minimum as shown in Fig. S-4(B). A small vesicle, when starting from the ridge quickly slides down and then moves along the axis maintaining its direction of motion (Fig. S-4(C)).

The corresponding trajectories are shown in second panel (Fig. S-4 (ii)). We also show the adhesion and bending energise for each case in Fig. S-4(iii-iv), that shows similar behaviour as for the other cases.

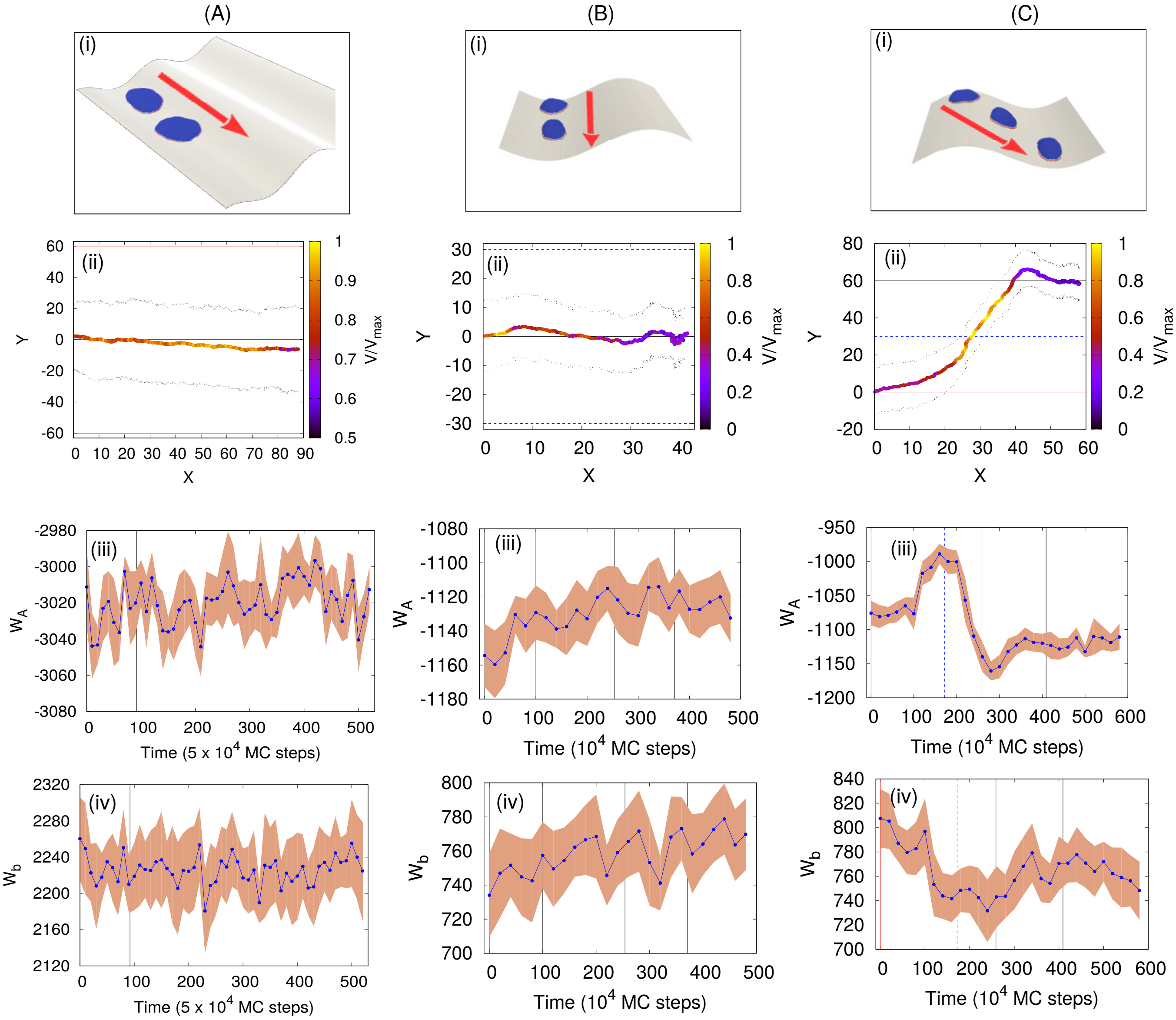


FIG. S-4. More trajectories for motile vesicle moving on a sinusoidal substrate with large wavelength. We use $z_m = 10\ l_{min}$; $y_m = 120\ l_{min}$. (A) A large vesicle starting from the minimum of the sinusoidal substrate continues its initial direction of migration. (B) A small vesicle starting from the minimum of the sinusoidal substrate continues to the minimum of the substrate. (C) Small vesicle starting from the maximum of a sinusoidal substrate shifts to the minimum. In (i), we show the configurations, in (ii), we show the trajectories with the speed of the vesicle in color code, in (iii), we show the adhesion energy and in (iv) , we show the bending energy. For large vesicle (A), we use $N = 3127$, $E_{ad} = 2.0$ $F = 1.0 k_B T/l_{min}$ and $\rho = 2.4\%$. For small vesicle (B-C), we use $N = 607$, $E_{ad} = 3.0\ k_B T$, $F = 2.0 k_B T/l_{min}$ and $\rho = 4.9\%$.

#### S-5. VARIATION OF PROTEIN-PROTEIN BINDING ENERGY FOR THE MIGRATION ON SINUSOIDAL SUBSTRATE WITH LARGE WAVELENGTH

Here, we show the variation of protein-protein binding energy with time when a large vesicle migrates on a sinusoidal substrate with $z_m = 10\ l_{min}$, $y_m = 120\ l_{min}$ (Fig. 2(C-D), main paper). We note that the energy is minimum when the vesicle is on the maximum of the substrate. This indicates that the proteins form much more strong cluster when the vesicle is on the maximum of the sinusoidal substrate. However, the variation in the protein-protein binding energy is much smaller in comparison with the other energies.

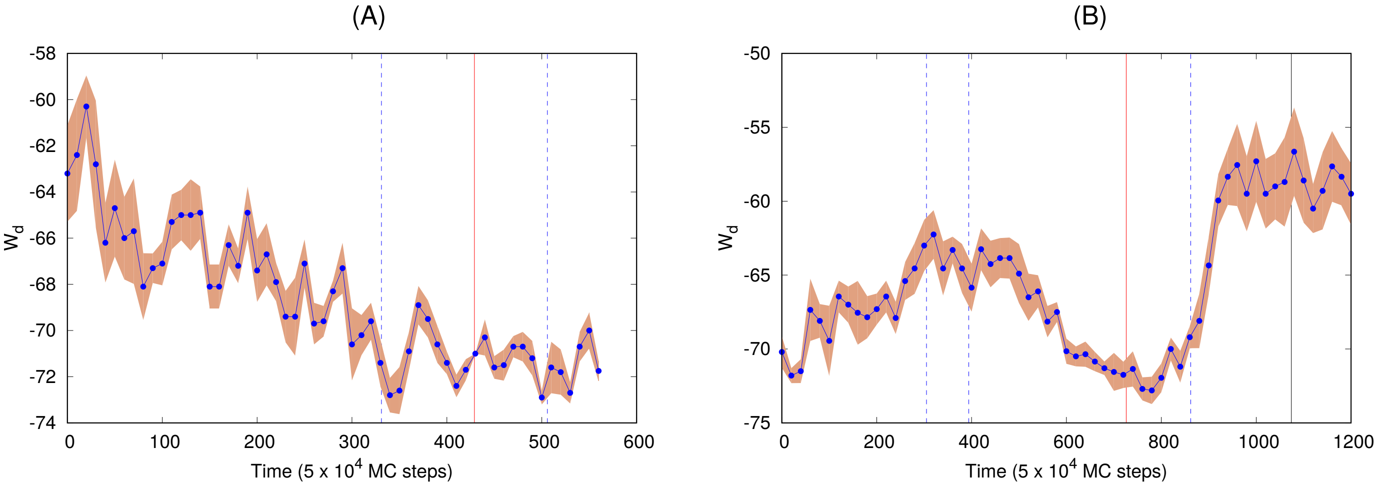


FIG. S-5. Variation of protein-protein binding energy when a large vesicle migrates on a sinusoidal substrate with $z_m = 10\ l_{min}$, $y_m = 120\ l_{min}$. For (A), we use $F = 2.0\ k_BT/l_{min}$ and for (B), we use $F = 1.0\ k_BT/l_{min}$. Other parameters are, $N = 3127$, $E_{ad} = 2.0\ k_BT$, $\rho = 2.4\ \%$

#### S-6. PERIODICITY IN THE SPEED FOR THE MIGRATION ON SINUSOIDAL SUBSTRATE WITH SMALL WAVELENGTH

In Fig. S-6, we show the distribution of speed in a full period of sinusoidal variation for the case of Fig.3 from the main text, for each of the four cases. Fig. S-6(A-B) shows nice periodicity for the simulation results, but Fig. S-6(C-D), does not show nice periodic variation from the migration of karatocytes, as the errors are larger in the case of experiment.

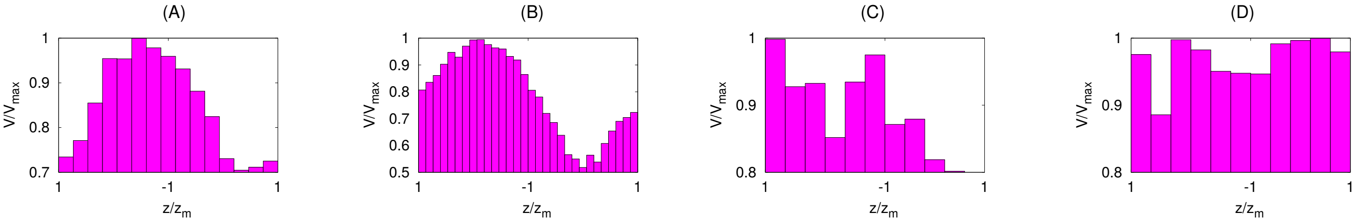


FIG. S-6. Periodicity in the speed for the migration on sinusoidal substrate with small wavelength (Fig. 3, main text). We plot the distribution of speed in a full period of sinusoidal variation for the case of Fig. 3, main text, for each of the four cases respectively. Here, $z/z_m = 1$ represents the ridges (maximum) while $z/z_m = -1$ represents the grooves (minimum) of the sinusoidal substrate. For simulation results, we use $N = 607$, $E_{ad} = 3.0\ k_BT$, $F = 4.0 k_BT/l_{min}$ and $\rho = 4.9\%$.

#### S-7. MORE TRAJECTORIES FOR THE MIGRATION ON SINUSOIDAL SUBSTRATE WITH SMALL WAVELENGTH

Here, we show few more trajectories for the sinusoidal substrate with small wavelength using the small vesicle only. In S-7(A-C), the vesicle does not maintain any particular direction and shows zig-zag like migration. In S-7(D) the vesicle finally migrates in a direction orthogonal to the sinusoidal substrate.

In second panel (S-7(ii)), we show the trajectories for each cases, and in the third panel (S-7(iii)) we show the speed showing oscillatory behaviour similar to the previous cases.

#### S-8. MORE TRAJECTORIES OF KARATOCYTES MIGRATING ON SINUSOIDAL SUBSTRATES

Here, we show the few more trajectories of the migration of karatocytes on sinusoidal substrate in Fig. S-8. In panel (i), we show the configurations, in second panel we show the corresponding trajectories and finally in the last

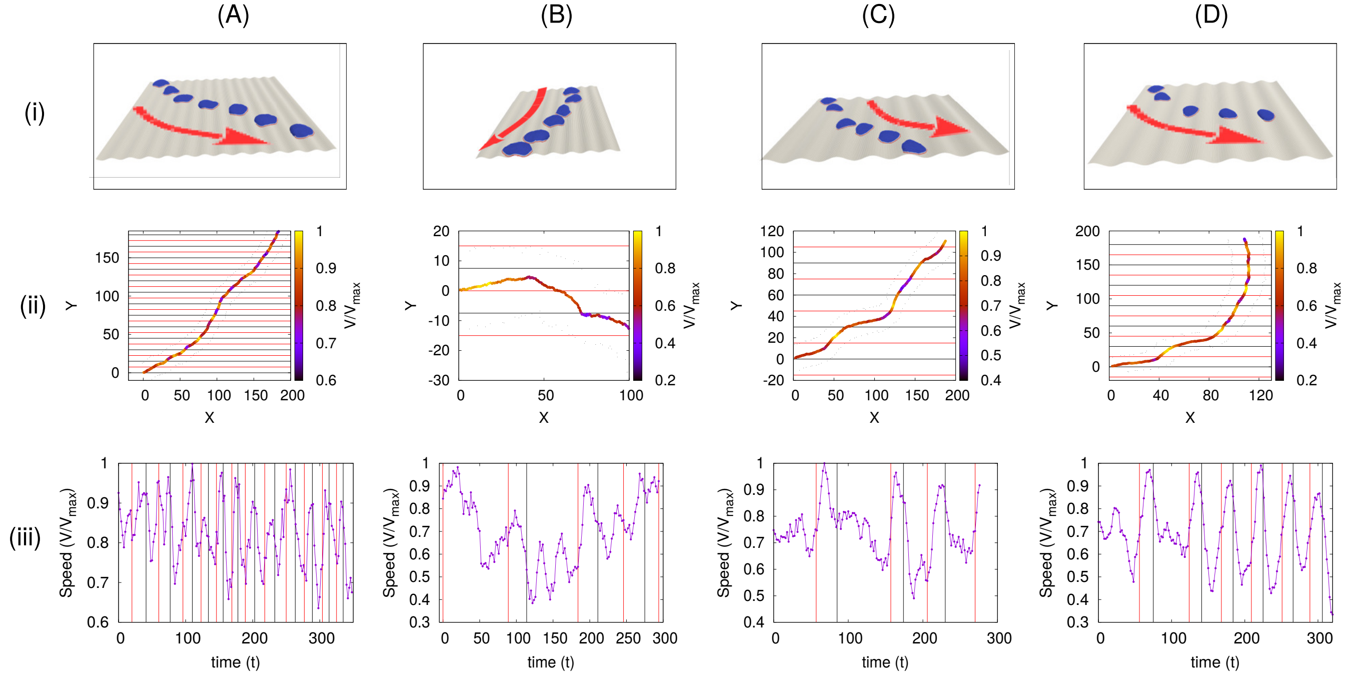


FIG. S-7. More trajectories for motile vesicle moving on a sinusoidal substrate with small wavelength. (A) A vesicle started from the minimum of the substrate shows zig-zag like motion with oscillating speed along the trajectory. (B) A vesicle started from the maximum of the sinusoidal substrate migrates with oscillating speed. (C-D) Two different simulations of vesicle starting from the minimum. For (A-B) we use $z_m = 1\ l_{min};\ y_m = 15\ l_{min}$ for the sinusoidal substrate, and for (C-D), we use $z_m = 2\ l_{min};\ y_m = 30\ l_{min}$. Other parameters are $N = 607$, $E_{ad} = 3.0\ k_BT$, $F = 4.0 k_BT/l_{min}$ and $\rho = 4.9\%$.

panel of Fig. S-8, we show the speed of the migrating cells with time, showing oscillatory behaviour.

#### S-9. VESICLE MIGRATING ON CYLINDRICAL FIBER: MAPPING FROM ‘TIME’ TO ‘MIGRATION ANGLE’

For the case of vesicle migrating on cylindrical fiber, we measure the quantities as a function of time, however, in MC simulations, the time is not a real quantity, rather it represents the MC time. So, we map time to a real quantity, i.e., the migration angle, which is defined as the angle between the displacement vector of the vesicle in a small time $\Delta t$ and the cylindrical axis. This quantity, initially starts from zero saturates to $\pi/2$ for larger times. We show this angle with time in Fig. S-9.

Since the measurement of angle ($\theta$) is associated with some statistical errors, we fit this data with a close quadratic function of the type $bt + ct^2$, that monotonically increases with time. This fitting allows us to describe different quantities as a function of migration angle, rather that unrealistic MC time.

#### S-10. VESICLE MIGRATING ON CYLINDRICAL FIBER: VARIOUS ENERGIES AS A FUNCTION OF TIME

In the main text (Fig. 4(B-D)), we show the variation of different energies of the vesicle as a function of its migration angle, when migrating on cylindrical fiber. Here, we present the same result as a function of MC time in Fig. S-10.

#### S-11. ENERGY OF AXIAL AND CIRCUMFERENTIAL ORIENTATION WITH FIBER RADIUS

Here, we show the total energy of the vesicle when it is in the axial orientation verses when it is in the circumferential orientation Fig.S-11(A). We note that the energy decreases with $R$, as the curvature of the cylinder decreases with $R$. Also, the difference between the energies in the axial and circumferential orientation also decreases with $R$.

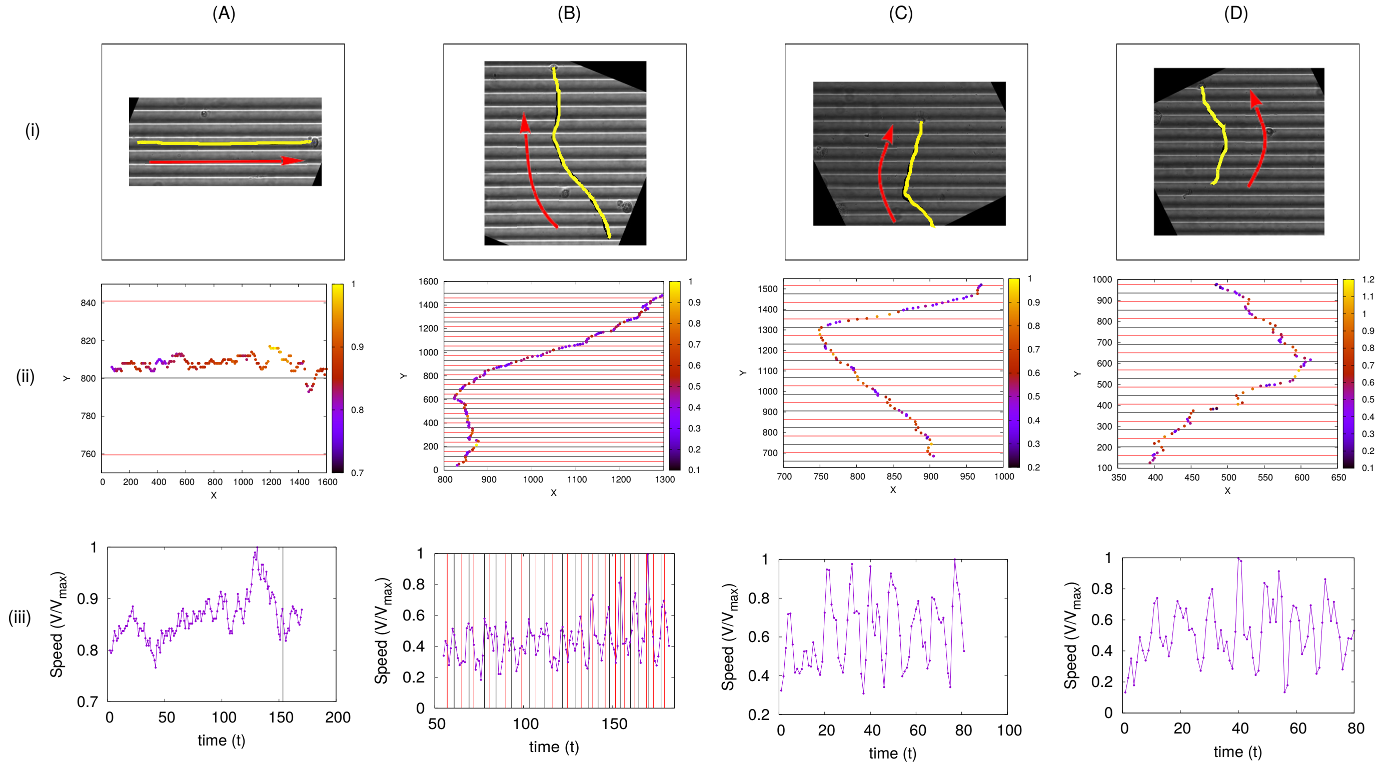


FIG. S-8. Here, we show few more trajectories for the migration of *karatocytes* on sinusoidal substrates. (A) The cell seems to move along the minimum without tending to change its direction of migration. (B-D) The cell migrates almost orthogonal to the sinusoidal substrate throughout its trajectories. Here, in (i) we show the snapshots, in (ii) we show the trajectories with the speed in the colour code, and in (iii), we show the speed of the cell with time, showing oscillatory behaviour.

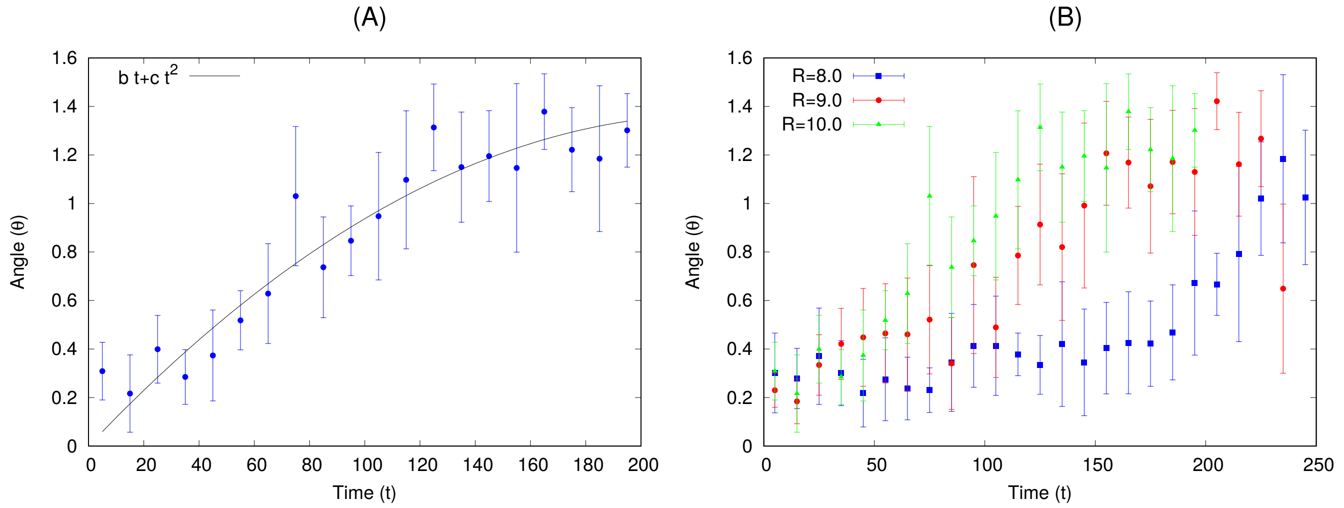


FIG. S-9. Migration angle with time for the case of vesicle migrating on a cylindrical fiber (Fig.4A). We define this quantity as the angle between the displacement vector of the vesicle in a small time $\Delta t$ and the cylindrical axis. (A) Variation of migration angle $(\theta)$ with time for $R = 10\ l_{min}$. We fit this data with a monotonic quadratic function of the type $bt + ct^2$, where, $b = 1.20085 \times 10^{-2}$ and $c = -2.63639 \times 10^{-5}$. (B) Migration ange $(\theta)$ for different values of $R$. The parameters are same as Fig. 4(A) of the main text.

In order to identify the anisotropy in the experimental data for D.d. cells migrating on fibers, we define the quantity called Curvotactic Anisotropy Parameter (CAP) as the absolute value of the velocity $|V_{||}|$ in the curved direction divided by the absolute value of the velocity in the perpendicular direction $|V_{\perp}|$,

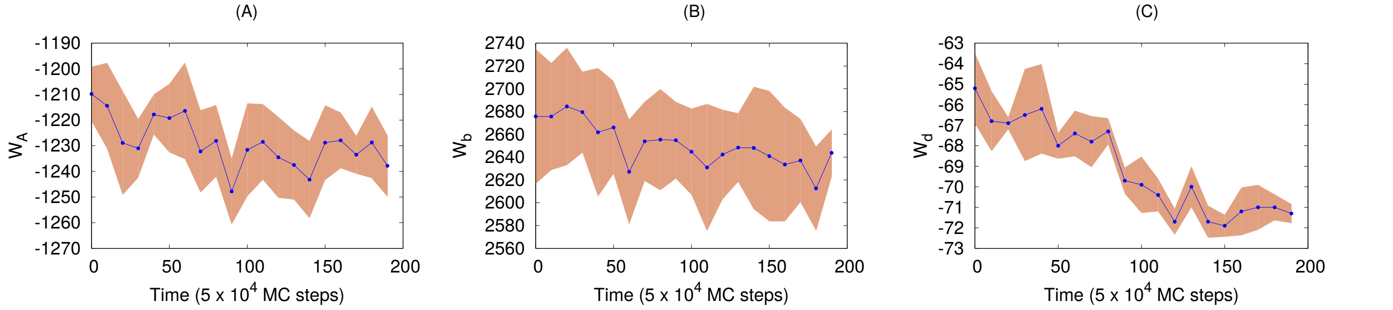


FIG. S-10. Different energies with time for a vesicle migrating on cylindrical fiber (Fig.4A). (A) Adhesion energy with time. (B) Bending energy with time. (C) Protein-protein binding energy with time. The parameters are same as Fig. 4(A) of the main text.

$$CAP = \frac{< |V_{||}| >}{< |V_{\perp}| >}$$

In Fig. S-11(B), we show this quantity for fibers of different radius of curvature. The CAP is found to be lower for larger radius of curvature, in agreement with the simulations that predict a smaller bias for circumferential orientation as the radius of the fiber increases (Fig. S-11A).

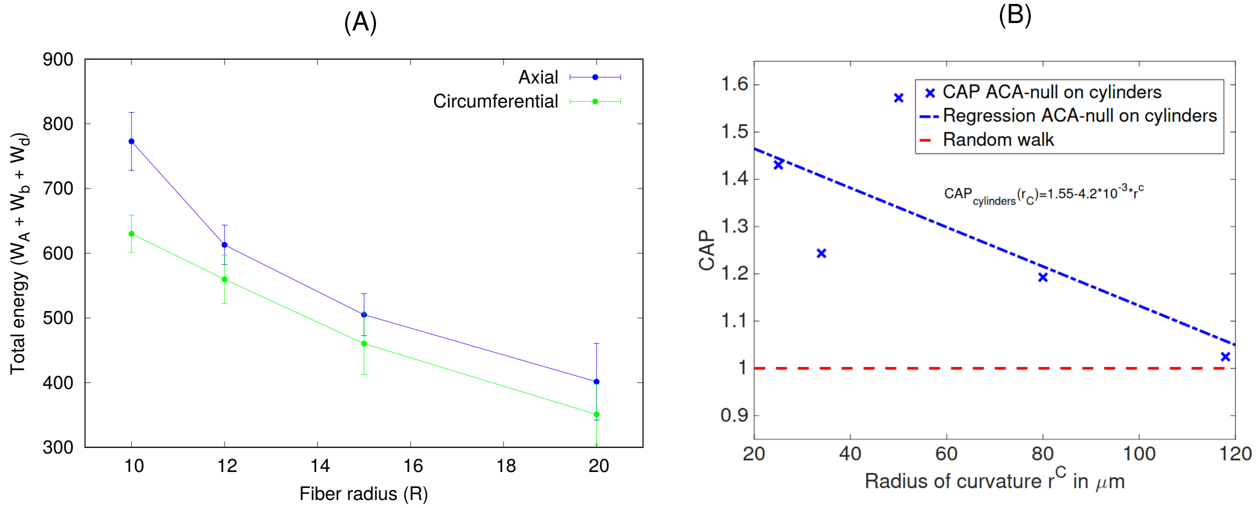


FIG. S-11. Quantification of the anisotropy of the migration of cells/vesicles on fibers of different radii. (A) The total energy of the vesicle as a function of radius of the fiber, comparing two cases: (1) When the vesicle is migrating along the axial direction of the fiber, and (2) When the vesicle is migrating along the circumferential direction. (B) The Curvotactic Anisotropy Parameter (CAP) as a function of radius of curvature of the fiber. For simulation results, the parameters are same as in Fig. 4A of the main text.

#### S-12. MDCK CELLS ON FIBER AND INSIDE TUBE

Here, we show the distribution of velocity of the migrating MDCK cells on the fiber (Fig. S-12(A)) and inside the tube (Fig. S-12(B)).

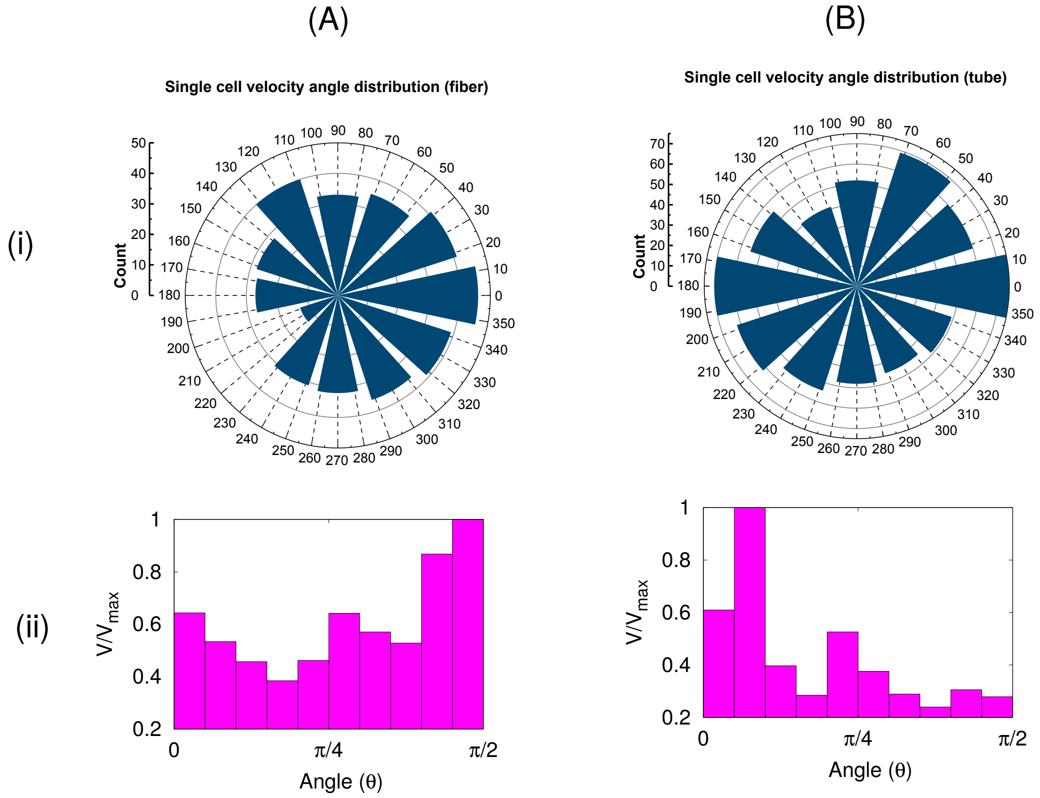


FIG. S-12. Angular speed distribution for MDCK cells migrating on (A) Fibers of 50 to 70$\mu m$ in diameter, and (B) Tubes of 67 to 75$\mu m$ in diameter. The direction $0° - 180°$ represents the circumferential direction and the direction along $90° - 270°$ is the axial direction.

#### S-13. VESICLE MIGRATING ON CYLINDER OF ELLIPTICAL CROSS-SECTION WITH DIFFERENT RATIO OF $R_x/R_y$

We note that the migrating vesicle, when exposed to a fiber of elliptical cross-section migrates and reorients in the circumferential direction when the aspect ratio $r = R_x/R_y$ is close to unity (Fig. S-13A). As the aspect ratio $r$ increases, the reorientation time also increases (Fig. S-13B). For a very large value of $r$, the vesicle only migrates along the axis and never rotates circumferentially (Fig. S-13C). We also show the angle of migration $\theta$ with time for three different values of $r$ in Fig. S-13D.

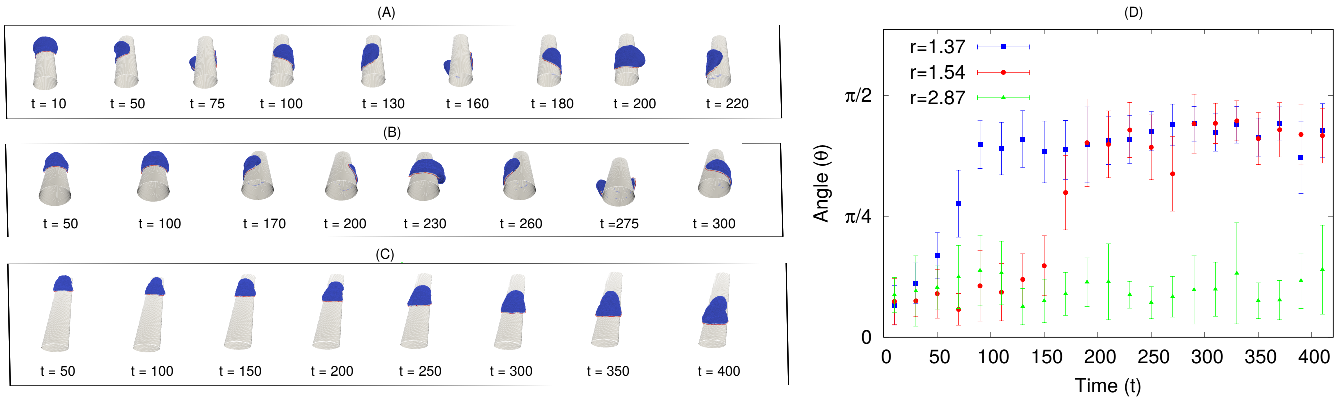


FIG. S-13. Vesicle migrating on elliptical fiber of different aspect ratio $r = R_x/R_y$ with fixed circumference $2\pi R$, with $R = 10\ l_{min}$. (A) Configuration with time for aspect ratio $r = 1.37$. (B) Configuration with time for aspect ratio $r = 1.54$. (C) Configuration with time for aspect ratio $r = 2.87$. (D) Angle of migration $\theta$ with time for a vesicle migrating on elliptical fiber with different values of $r$. Other parameters are $E_{ad} = 1.5\ k_BT$, $F = 2.0\ k_BT/l_{min}$, and $\rho = 2.4$ %.

#### S-14. SPREADING AND MIGRATION OF VESICLE INSIDE A CYLINDRICAL TUBE

Here, we show the results for the (non-motile) vesicle spreading inside a cylindrical tube (Fig. S-14). The vesicle seems to elongate in the axial direction without any preference to orient circumferentially.

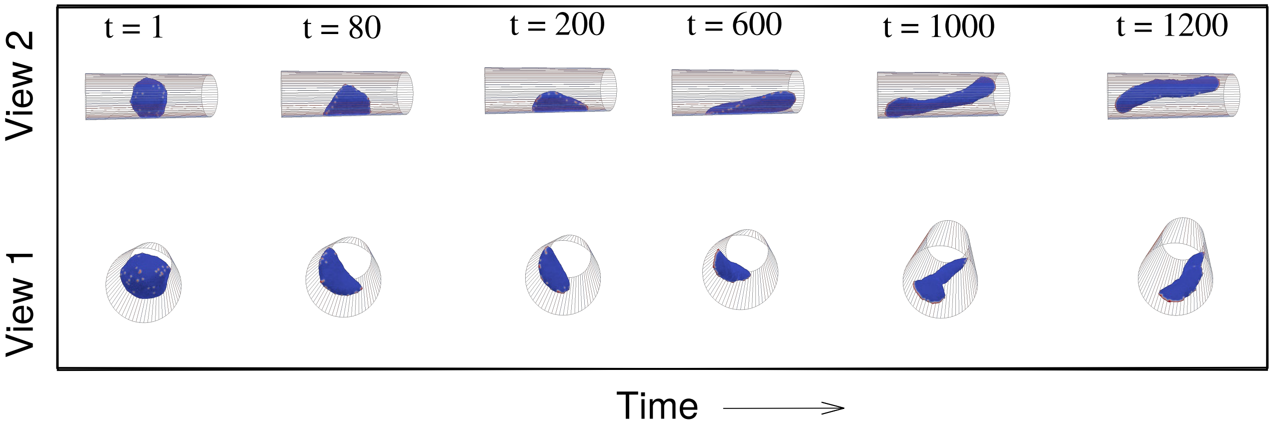


FIG. S-14. Vesicle inside a cylinder: two arc formation. We show the configurations of a vesicle inside a cylinder, starting from a quasi-spherical vesicle. Here, we use $R = 23$. Other parameter values are same as Fig. 6A of the main text.

Next, we study the migration of cells inside a cylindrical tube, a uniformly concave surface. We start with our vesicle, that shows no tendency to rotate circumferentially when initially aligned to migrate along the axial direction (Fig.S-15A, Movie-S29). Over time, the vesicle is found to lose its motility, and the leading edge protein cluster breaks into several parts, which leads to a decrease in the total active force that propels the vesicle (Fig.S-15B). This is similar to the behavior inside the grooves of the sinusoidal surface (Fig.2A,B, main text). We chose here a tube radius such that the circumference of the tube is much larger than the vesicle's diameter. In this regime we can explore the cell migration on the surface, avoiding "plugging" of the tube by the vesicle when the tube radius is smaller than the cell radius [23].

The different energy terms of the vesicle do not show any systematic variation during its migration in the tube (Fig.S-15C-E). In Fig.S-15F we show that all the proteins along the cell edge are well adhered to the substrate, which is why the leading edge can easily break up and form clusters along any direction. The leading-edge that was initially oriented axially, will tend to break into two arcs that point side-ways. This destabilizes the polarized leading-edge aggregate, and the cell migration slows down, sometimes to a halt (forming a two-arc non-motile phenotype). As a result, we find that the vesicle in the tube ends up either as slowly migrating in the axial direction (on average, Fig.S-15A) or a two-arc (non-motile) vesicle that can be axial or "bridge" orthogonally to the axis (Fig. S-14, Movie-S30).

Comparing to experimental observations of cells moving inside tubes [27], it was indeed found that cells tend to migrate along the tube axis, as we obtain (Fig.S-15A). However, this tendency is strongly cell-type dependent, with some cells becoming non-motile inside tubes, forming adhesion "bridges" that can be orthogonal to the tube axis [28]. This observation agrees with our finding that the motility inside the tube is strongly inhibited, with the cells tending to lose their polarization (Fig.S-15A,B). In [23], it was indeed observed that the motility along the tube axis decreases as the tube radius decreases, as cells migrate less persistently, and their leading-edge lamellipodia becomes less persistent and less stable. In addition, the overall orientation of actin filaments for cells inside tubes was axial, in qualitative agreement with our model's results.

Fig.S-15G, we show the trajectories for the migration of MDCK cells on the inside of a tube. The cells were found to be weakly motile, with the most persistent motility periods aligned with the tube axis (Movie-S31), in agreement with the model predictions.

Another recent study [29] on two cell types (endothelial and epithelial cells), found similar axial alignment of the cells shape and their migration inside tubes. However, the role of stress-fibers, which we do not include in our model, was suggested to have a major role for these cells.

In Fig. S-16 we summarize the steady-state shapes for the simulated motile vesicles on the different curved substrates. We note that the adhered area is maximal for a flat substrate, but is not very different when the vesicle is inside the tube, where it is also well adhered. When on the fiber, moving in the axial direction, the adhered area is the smallest, and it is slightly larger when moving in the circumferential direction. The aspect ratio shows that the active force is able to stretch the vesicle sideways along the axis of the tube, when it is oriented circumferentially.

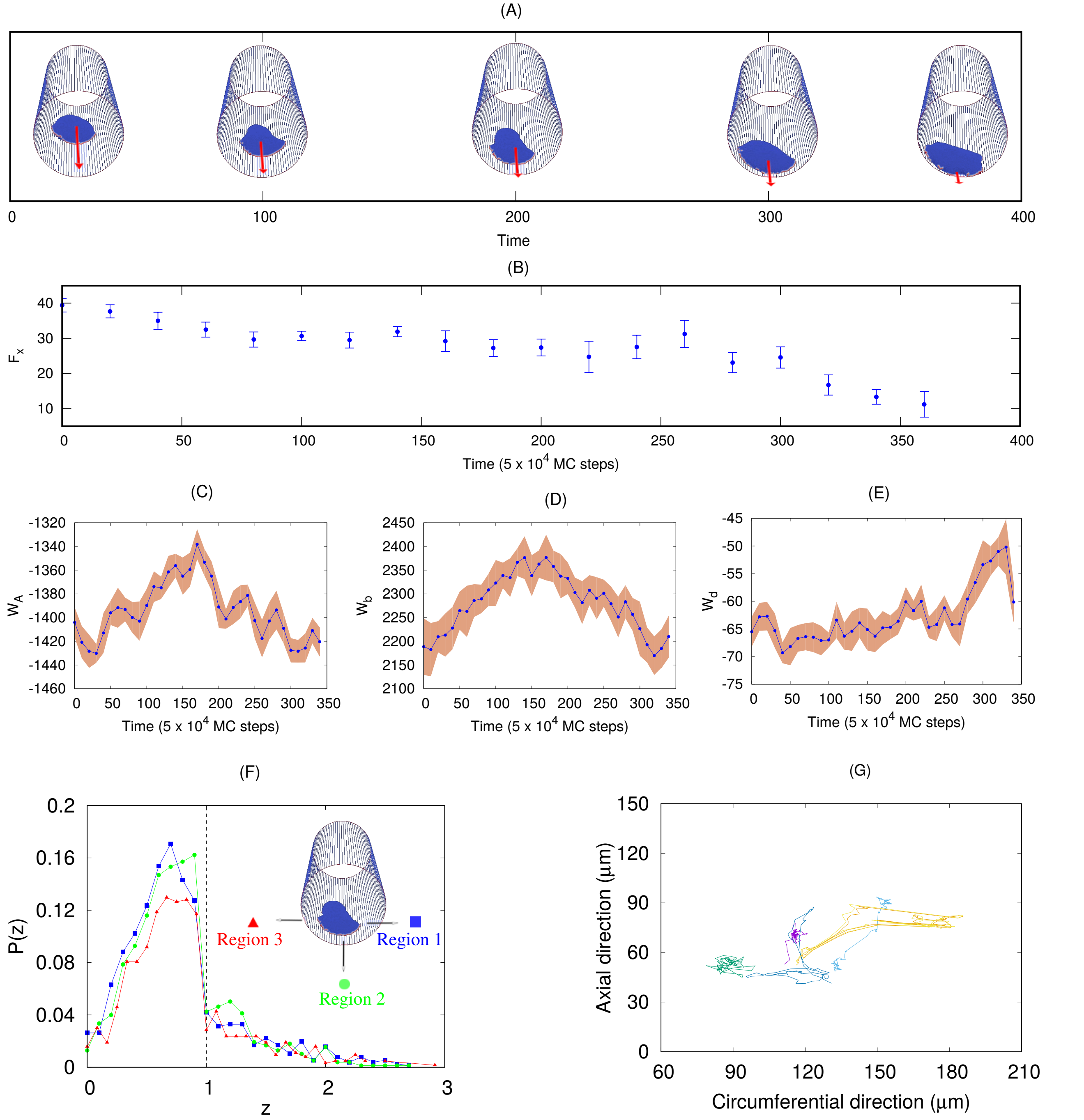


FIG. S-15. Vesicle migrating inside a cylindrical tube. (A) Configuration of vesicle migrating inside tube, initiated in the axial direction. (B) Magnitude of active force along the axis of cylinder ($F_x$) with time. (C) The adhesion energy of the vesicle with time. (D) The bending energy of the vesicle with time. (E) The binding energy between proteins with time. (F) Probability distribution of a protein at $z$ distance above the cylindrical substrate. The left side of the vertical dashed line (at $z = 1$) represents adhered proteins. Here, we use $R = 35\ l_{min}$, $E_{ad} = 1.0$, $F = 2.0$, and $\rho = 2.4\%$. (G) Trajectories of the MDCK cells migrating inside tubes of $67 - 75\ \mu m$ in diameter.

The highly elongated shape of the aligned cell moving in the tube, compared to the flat substrate, fits well with the observed elongation of the cells when migrating inside the grooves of the sinusoidal substrate [30].

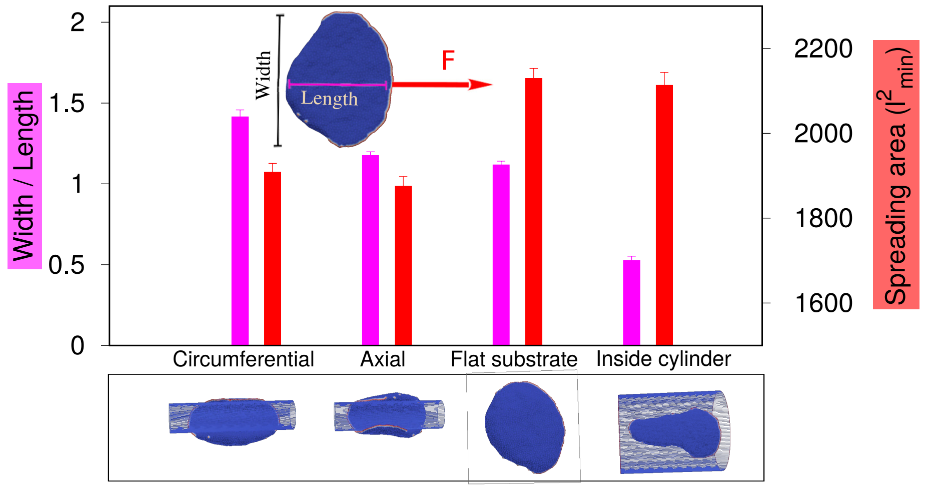


FIG. S-16. Ratio of the length (along $F$) and width (orthogonal to $F$) of the vesicle and the adhered area, for four different cases when it is moving (1) along circumferential direction on a cylinder, (2) along axial direction on a cylinder, (3) on a flat substrate and (4) inside of a cylindrical tube. The magenta color is for the aspect ratio and the red color showing the adhered area. Inset shows a crescent shape on a flat substrate, defining the length and the width of the vesicle. For fiber, we use $R = 10\ l_{min}$, for tube, we use $R = 21\ l_{min}$. Other parameters are $F = 2.0$, $E_{ad} = 1.0$ and $\rho = 2.4\%$.

#### SUPPLEMENTARY MOVIES

High resolution movies are also available here.

- **Movie-S1** (Fig. S-1) Preparation of vesicle on curved substrate. For sinusoidal substrate, we vary $z_m$ from $1 - 10\ l_{min}$ while keeping $y_m$ fixed ($= 120\ l_{min}$). The other parameters are: $N = 3127$, $E_{ad} = 2.0\ k_BT$, $F = 2.0\ k_BT/l_{min}$, and $\rho = 2.4$ %.

- **Movie-S2** (Fig. 2A) Small sized vesicle (N=607) migrating along the axis of sinusoidal substrate starting from the minimum (groove) of the substrate. For sinusoidal substrate, we use $z_m = 10\ l_{min}; y_m = 120\ l_{min}$. The other parameters are: $E_{ad} = 3.0\ k_BT$, $F = 4.0\ k_BT/l_{min}$, and $\rho = 4.9$ %.

- **Movie-S3** (Fig. 2B) Small sized vesicle (N=607) migrating along the axis of sinusoidal substrate starting from the maximum (ridge) of the substrate. For sinusoidal substrate, we use $z_m = 10\ l_{min}; y_m = 120\ l_{min}$. The other parameters are: $E_{ad} = 3.0\ k_BT$, $F = 4.0\ k_BT/l_{min}$, and $\rho = 4.9$ %.

- **Movie-S4** (Fig. 2C) Large sized vesicle (N=3127) migrating along the axis of sinusoidal substrate starting from the minimum of the substrate. For sinusoidal substrate, we use $z_m = 10\ l_{min}; y_m = 120\ l_{min}$. The other parameters are: $E_{ad} = 2.0\ k_BT$, $F = 2.0\ k_BT/l_{min}$, and $\rho = 2.4$ %.

- **Movie-S5** (Fig. 2D) Large sized vesicle (N=3127) migrating along the axis of sinusoidal substrate starting from the maximum of the substrate. For sinusoidal substrate, we use $z_m = 10\ l_{min}; y_m = 120\ l_{min}$. The other parameters are: $E_{ad} = 2.0\ k_BT$, $F = 1.0\ k_BT/l_{min}$, and $\rho = 2.4$ %.

- **Movie-S6** (Fig. S-2A) Large sized vesicle (N=3127) migrating along the axis of sinusoidal substrate starting from the minimum of the substrate. For sinusoidal substrate, we use $z_m = 10\ l_{min}; y_m = 120\ l_{min}$. The other parameters are: $E_{ad} = 2.0\ k_BT$, $F = 1.0\ k_BT/l_{min}$, and $\rho = 2.4$ %.

- **Movie-S7** (Fig. S-2B) Small sized vesicle (N=607) migrating along the axis of sinusoidal substrate starting from the minimum of the substrate. For sinusoidal substrate, we use $z_m = 10\ l_{min}; y_m = 120\ l_{min}$. The other parameters are: $E_{ad} = 3.0\ k_BT$, $F = 2.0\ k_BT/l_{min}$, and $\rho = 4.9$ %.

- **Movie-S8** (Fig. S-2C) Small sized vesicle (N=607) migrating along the axis of sinusoidal substrate starting from the maximum of the substrate. For sinusoidal substrate, we use $z_m = 10\ l_{min}; y_m = 120\ l_{min}$. The other parameters are: $E_{ad} = 3.0\ k_BT$, $F = 2.0\ k_BT/l_{min}$, and $\rho = 4.9$ %.

- **Movie-S9** (Fig. 3A) Small sized vesicle (N=607) migrating orthogonal to the axis of sinusoidal substrate with small wavelength. For sinusoidal substrate, we use $z_m = 1\ l_{min}; y_m = 15\ l_{min}$. The other parameters are: $E_{ad} = 3.0\ k_BT$, $F = 4.0\ k_BT/l_{min}$, and $\rho = 4.9$ %.
- **Movie-S10** (Fig. 3B) Small sized vesicle (N=607) initiating from the maximum of the sinusoidal substrate of small wavelength finally becomes orthogonal to the sinusoidal axis. For sinusoidal substrate, we use $z_m = 2\ l_{min}; y_m = 30\ l_{min}$. The other parameters are: $E_{ad} = 3.0\ k_BT$, $F = 4.0\ k_BT/l_{min}$, and $\rho = 4.9$ %.
- **Movie-S11** (Fig. S-4A) Small sized vesicle (N=607) migrating on sinusoidal substrate at an angle $\sim\ 45°$ to the sinusoidal axis. For sinusoidal substrate, we use $z_m = 1\ l_{min}; y_m = 15\ l_{min}$. The other parameters are: $E_{ad} = 3.0\ k_BT$, $F = 4.0\ k_BT/l_{min}$, and $\rho = 4.9$ %.
- **Movie-S12** (Fig. S-4B) Small sized vesicle (N=607) migrating on sinusoidal substrate along the sinusoidal axis. For sinusoidal substrate, we use $z_m = 1\ l_{min}; y_m = 15\ l_{min}$. The other parameters are: $E_{ad} = 3.0\ k_BT$, $F = 4.0\ k_BT/l_{min}$, and $\rho = 4.9$ %.
- **Movie-S13** (Fig. S-4C) Small sized vesicle (N=607) migrating on sinusoidal substrate initially along the sinusoidal axis finally migrates at an angle $\sim\ 45°$. For sinusoidal substrate, we use $z_m = 2\ l_{min}; y_m = 30\ l_{min}$. The other parameters are: $E_{ad} = 3.0\ k_BT$, $F = 4.0\ k_BT/l_{min}$, and $\rho = 4.9$ %.
- **Movie-S14** (Fig. S-4D) Small sized vesicle (N=607) migrating on sinusoidal substrate initially along the sinusoidal axis finally becomes orthogonal to the sinusoidal axis. For sinusoidal substrate, we use $z_m = 2\ l_{min}; y_m = 30\ l_{min}$. The other parameters are: $E_{ad} = 3.0\ k_BT$, $F = 4.0\ k_BT/l_{min}$, and $\rho = 4.9$ %.
- **Movie-S15** (Fig. 3C) Karatocytes migrating on sinusoidal substrate at an angle.
- **Movie-S16** (Fig. 3D) Karatocytes migrating on sinusoidal substrate, initially along the axis changes its direction.
- **Movie-S17** (Fig. S-5A) Karatocytes migrating on sinusoidal substrate along the axis maintains its direction of migration.
- **Movie-S18** (Fig. S-5B) Karatocytes migrating on sinusoidal substrate in the orthogonal direction to the sinusoidal axis.
- **Movie-S19** (Fig. S-5C) Karatocytes migrating on sinusoidal substrate almost orthogonal direction to the sinusoidal axis.
- **Movie-S20** (Fig. S-5D) Karatocytes migrating on sinusoidal substrate closely orthogonal direction to the sinusoidal axis.
- **Movie-S21** (Fig. 4A) Vesicle migrating along the axis of a cylindrical fiber, finally migrates in the circumferential direction. Parameters are: $R = 10\ l_{min}$, $E_{ad} = 1.0\ k_BT$, $F = 2.0\ k_BT/l_{min}$, and $\rho = 2.4$ %.
- **Movie-S22** (Fig. 4G) Overlay of the first micrograph of a time series of migrating cells and the corresponding 43 trajectories extracted from this series. We used AX2 cells labeled with LimE-GFP. The diameter of the optical fiber is $160\mu m$ and the images recorded with an Olympus confocal laser scanning microscope Fluoview 1000 in the DIC mode. The different colours of the trajectories are used to distinguish between individual trajectories. Gray scale represents the DIC intensity in a.u., while the black scale bar corresponds $100\mu m$.
- **Movie-S23** (Fig. 4J) $2D$ time-lapse projection of a MDCK-LifeAct-GFP cell on a microfiber of $50\mu m$ in diameter, showing preferential migration along the circumferential axis of the fiber. L-axis and C-axis indicate the cylindrical longitudinal axis and circumferential axis, respectively. Magenta line indicates the cell trajectory.
- **Movie-S24** (Fig. 5A, top panel) D.d.. cells migrating on micropillars of circular cross-section.
- **Movie-S25** (Fig. 5A, bottom panel) D.d.. cells migrating on micropillars of triangular cross-section.
- **Movie-S26** (Fig. 5F) Vesicle migrating on an elliptical cylinder. Parameters are: $R_x = 12\ l_{min}$, $R_y = 7.773\ l_{min}$, $E_{ad} = 1.5\ k_BT$, $F = 2.0\ k_BT/l_{min}$, and $\rho = 2.4$ %.
- **Movie-S27** (Fig. S-9A) Vesicle migrating on an elliptical cylinder. Parameters are: $R_x = 11.5\ l_{min}$, $R_y = 8.377\ l_{min}$, $E_{ad} = 1.5\ k_BT$, $F = 2.0\ k_BT/l_{min}$, and $\rho = 2.4$ %.

- **Movie-S28** (Fig. S-9C) Vesicle migrating on an elliptical cylinder. Parameters are: $R_x = 14\ l_{min}$, $R_y = 4.882\ l_{min}$, $E_{ad} = 1.5\ k_BT$, $F = 2.0\ k_BT/l_{min}$, and $\rho = 2.4$ %.
- **Movie-S29** (Fig. 6A) Vesicle migrating inside a cylindrical tube. Parameters are: $R = 35\ l_{min}$, $E_{ad} = 1.0\ k_BT$, $F = 2.0\ k_BT/l_{min}$, and $\rho = 2.4$ %.
- **Movie-S30** (Fig. S-11) Non-migrating vesicle spreading and elongating inside a cylindrical tube. Parameters are: $R = 23\ l_{min}$, $E_{ad} = 1.0\ k_BT$, $F = 2.0\ k_BT/l_{min}$, and $\rho = 2.4$ %.
- **Movie-S31** (Fig. 6G) $2D$ time-lapse projection of a MDCK-LifeAct-GFP cell in a microtube of 65 $\mu m$ in diameter, showing preferential migration along the longitudinal axis of the tube. L-axis and C-axis indicate the cylindrical longitudinal axis and circumferential axis, respectively. Magenta line indicates the cell trajectory.

---


[1] N. Ramakrishnan, P.B. Sunil Kumar, and Ravi Radhakrishnan. Mesoscale computational studies of membrane bilayer remodeling by curvature-inducing proteins. *Physics Reports*, 543(1):1–60, 2014. Mesoscale computational studies of membrane bilayer remodeling by curvature-inducing proteins.

[2] Raj Kumar Sadhu, Samo Penič, Aleš Iglič, and Nir S. Gov. Modelling cellular spreading and emergence of motility in the presence of curved membrane proteins and active cytoskeleton forces. *The European Physical Journal Plus*, 136(5):495, May 2021.

[3] Miha Fošnarič, Samo Penič, Aleš Iglič, Veronika Kralj-Iglič, Mitja Drab, and Nir S. Gov. Theoretical study of vesicle shapes driven by coupling curved proteins and active cytoskeletal forces. *Soft Matter*, 15:5319–5330, 2019.

[4] Raj Kumar Sadhu, Christian Hernandez-Padilla, Yael Eshed Eisenbach, Lixia Zhang, Harshad D Vishwasrao, Bahareh Behkam, Hari Shroff, Aleš Iglič, Elior Peles, Amrinder S. Nain, and Nir S Gov. Coiling of cellular protrusions around extracellular fibers. *Arxiv*, 2205.12488, 2022.

[5] Raj Kumar Sadhu, Sarah R Barger, Samo Penič, Aleš Iglič, Mira Krendel, Nils Gauthier, and Nir Gov. Theoretical model of efficient phagocytosis driven by curved membrane proteins and active cytoskeleton forces. *Soft Matter*, pages –, 2022.

[21] Christoph Blum. Curvotaxis and pattern formation in the actin cortex of motile cells. 2015.

[22] Marcel Schröder. Cell-substrate adhesion and contact guidance of dictyostelium discoideum on surfaces of varying curvatures. Master's thesis, Georg August University of Göttingen, 2018.

[23] Wang Xi, Surabhi Sonam, Thuan Beng Saw, Benoit Ladoux, and Chwee Teck Lim. Emergent patterns of collective cell migration under tubular confinement. *Nature communications*, 8(1):1–15, 2017.

[24] Alexandros Glentis, Carles Blanch-Mercader, Lakshmi Balasubramaniam, Thuan Beng Saw, Joseph d'Alessandro, Sebastien Janel, Audrey Douanier, Benedicte Delaval, Frank Lafont, Chwee Teck Lim, Delphine Delacour, Jacques Prost, Wang Xi, and Benoit Ladoux. The emergence of spontaneous coordinated epithelial rotation on cylindrical curved surfaces. *Science Advances*, 8(37):eabn5406, 2022.

[25] Gareth Bloomfield, David Traynor, Sophia P Sander, Douwe M Veltman, Justin A Pachebat, and Robert R Kay. Neurofibromin controls macropinocytosis and phagocytosis in *Dictyostelium*. *eLife*, 4:e04940, mar 2015.

[26] Sven Flemming, Francesc Font, Sergio Alonso, and Carsten Beta. How cortical waves drive fission of motile cells. *Proceedings of the National Academy of Sciences*, 117(12):6330–6338, 2020.

[27] Cas van der Putten, Daniëlle van den Broek, and Nicholas A Kurniawan. Myofibroblast transdifferentiation of keratocytes results in slower migration and lower sensitivity to mesoscale curvatures. *Frontiers in cell and developmental biology*, 10, 2022.

[28] Maike Werner, Ansgar Petersen, Nicholas A. Kurniawan, and Carlijn V. C. Bouten. Cell-perceived substrate curvature dynamically coordinates the direction, speed, and persistence of stromal cell migration. *Advanced Biosystems*, 3(10):1900080, 2019.

[29] Xiaoyu Yu, Haiqin Wang, Fangfu Ye, Xiaochen Wang, Qihui Fan, and Xinpeng Xu. Biphasic curvature-dependence of cell migration inside microcylinders: persistent randomness versus directionality. *bioRxiv*, 2022.

[30] Kwang Hoon Song, Sung Jea Park, Dong Sung Kim, and Junsang Doh. Sinusoidal wavy surfaces for curvature-guided migration of t lymphocytes. *Biomaterials*, 51:151–160, 2015.
